# Clustered structural variant hotspots enable oncogenic addiction and plasticity in osteosarcoma

**DOI:** 10.64898/2026.09.14.751472

**Authors:** Shreya Raman, Thorsten Kaltenbacher, David Merrell, Kavya Prasad, Yutaro Tanaka, Jakob Weiss, Emily Iannetta, Byron Butaney, Emily Jorswieck, Amr Salem, Lorena Lazo De La Vega, Evelina Ceca, Julia M.W. Wong, Suzanne J. Forrest, Kelly Klega, Mohammad Tanhaemami, Cora A. Ricker, Jinyu Wang, Diane M. Diehl, Jeremy Johnson, Carrie Cibulskis, Kristy Schlueter-Kuck, Liam Q. Alley, Natalie B. Collins, Cheng-Zhong Zhang, David S. Shulman, Brian D. Crompton, Katherine A. Janeway, Gad Getz, Riaz Gillani

## Abstract

The genomic landscape of osteosarcoma, the most common bone cancer worldwide, is among the most structurally complex of all human malignancies. The identification of recurrent and functionally consequential patterns has thus remained a challenge. Across 236 whole-genome sequencing osteosarcoma samples, we uncovered five genomic hotspots of clustered structural variation collectively altered in 58% of tumors. Four were associated with amplification of oncogenes (*MYC*, *CCND3*, *CCNE1*, *CDK4)*, while the fifth mapped largely upstream of *TP53*. Hotspot events showed coordinated patterns of co-occurrence and mutual exclusivity with each other and with tumor suppressor alterations, suggesting genomic context-specific selection. We found localized transcriptional dysregulation at hotspot event loci, and single cells harboring these events converged on a neural crest-like program, linking these structural alterations to a less differentiated cell state. These events were also detectable non-invasively through liquid biopsies and displayed ongoing structural evolution throughout disease progression, a finding with potential clinical utility. Our results provide novel insight into how complex rearrangements shape oncogenesis in osteosarcoma, with broader relevance to other cancers characterized by complex genomes.

## Introduction

Osteosarcoma is the most common primary bone malignancy, with half of all cases occurring in children and young adults^1^. Patients with localized osteosarcoma have a 5-year survival rate of ∼60%, which declines to ∼20% among those with metastatic or recurrent disease^2,3^. Approximately 55% of patients fall into this poor-prognosis category, comprising the ∼15% who present with metastasis at diagnosis and the ∼40% who present with localized disease but progress or relapse following treatment^2,3^. Current treatment regimens involve surgical resection and systemic multiagent chemotherapy, with no improvement in treatment options or outcomes seen in over four decades^4^.

One of the reasons for this therapeutic roadblock is the extensive heterogeneity across tumors which complicates the development of targeted therapies. Osteosarcomas show near ubiquitous loss of *TP53*, which enables genomic instability^5^. This instability manifests as a wide range of structural and numerical chromosomal abnormalities that contribute to the disease’s complexity and aggressiveness. In fact, osteosarcoma carries the highest burden of structural variants (SVs) among pediatric cancers^6^. A major mutational process driving these genomic rearrangements is chromothripsis, which involves the shattering of chromosomes followed by re-ligation of fragments in a random order. Loss-translocation-amplification (LTA) chromothripsis is one such recently described mechanism that enables the simultaneous inactivation of tumor suppressors, most frequently *TP53*, and amplification of oncogenes that can span multiple chromosomes via breakage-fusion-bridge cycles^7^.

Such foundational studies have expanded our understanding of the unifying molecular characteristics and recurrent mechanisms of tumor evolution in osteosarcoma^6–9^. Despite this progress, an understanding of complex genomic rearrangement patterns across osteosarcoma tumors and their functional consequences remains elusive. To address this gap, we assembled and analyzed whole genome sequencing (WGS) data from a cohort of 236 tumor-normal matched osteosarcoma samples to systematically dissect the role of complex rearrangement events in oncogenesis. We leveraged paired bulk and single-cell RNA-seq data to resolve the transcriptional changes and cellular states associated with these genomic events. Lastly, we demonstrated that these events could be detected and longitudinally monitored via non-invasive liquid biopsy. Together, these analyses represent a meaningful step forward in uncovering biologically and clinically relevant patterns within the apparent disorder of the osteosarcoma genome.

## Results

### Analysis of SV breakpoints in osteosarcoma tumors reveals hotspots of clustered SVs

We assembled and uniformly processed a WGS dataset of 236 tumor-normal paired osteosarcoma samples from 207 unique patients (**Fig. 1a, Extended Data Fig. 1a**). For 21 patients, we analyzed multiple longitudinally or spatially distinct tumor samples, encompassing multi-region diagnostic specimens as well as metastatic and relapse tumor tissues. The dataset additionally included 120 paired bulk RNA-seq and 14 paired single nucleus RNA-seq (snRNA-seq) samples (**Fig. 1b, Supplementary Table 1**). While most of this cohort was derived from previously published studies^10–13^, we generated WGS data for 19 samples and matched bulk RNA-seq data for 8 of these cases, 6 of which also had recently published snRNA-seq data available^14^.

**Figure 1:**
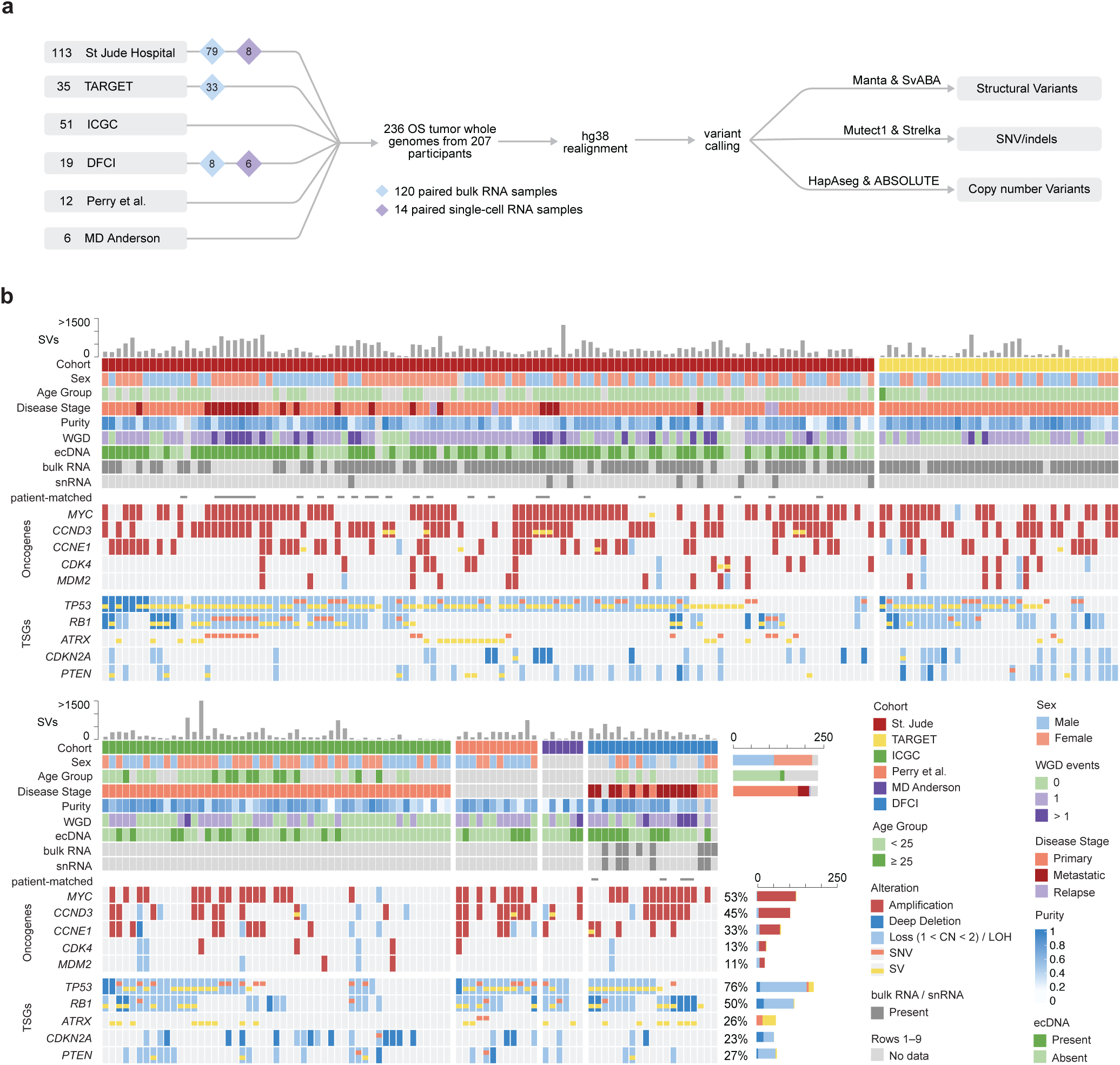
Study design and cohort overview. **a,** Overview of the osteosarcoma cohort comprising 236 tumor whole genomes from 207 participants across six cohorts: St Jude Hospital (n = 113), TARGET (n = 35), ICGC (n = 51), DFCI (n = 19), Perry et al. (n = 12) and MD Anderson (n = 6). Matched bulk RNA-seq (n = 120) and single-cell RNA-seq (n = 14) samples are indicated by blue and purple diamonds, respectively. Whole genomes were realigned to hg38 and analyzed for structural variants, SNVs/indels, and copy-number variants. **b,** Genomic and clinical landscape of the osteosarcoma cohort. Each column represents a tumor sample, with samples from the same patient placed adjacently and marked by horizontal bars. The top bar plot shows the total structural variant (SV) burden per sample. Tracks below display clinical characteristics, tumor purity, whole-genome doubling (WGD) status, ecDNA status, availability of matched bulk RNA-seq and snRNA-seq data, and somatic alterations affecting tumor suppressor genes (TSG) and oncogenic drivers. Copy number data was not available for *ATRX*. Plots on the right show the frequencies of clinical features and genomic alterations across the cohort. For genomic alterations, percentages indicate the proportion of samples with any alteration in each gene while the bars show the number of samples by alteration type. Samples with multiple alteration types in the same gene were assigned to a single alteration category using the following priority: deletion > LOH > amplification > SNV > SV.

We comprehensively characterized the mutational landscape by performing genome-wide identification of somatic SVs, single nucleotide variants (SNVs), and copy number alterations (SCNAs). We called SVs using two algorithms, Manta^15^ and SvABA^16^, and took the union of calls (64.5% concordance between callers) to maximize sensitivity (**Extended Data Fig. 1b**).

To assemble a non-redundant cohort, we selected one tumor genome per patient, prioritizing primary over metastatic or relapsed samples and, where multiple samples remained, the highest-purity sample. Across these 207 unique patient genomes, we observed a median of 215 SVs per tumor (IQR: 90–328). To characterize the landscape of structural variation, we first searched for genomic regions enriched with SV breakpoints. Following a previously described approach^17^, we quantified the number of samples with breakpoints across 50kb bins and identified those exceeding a background rate modeled by replication timing, GC content, mappability, heterochromatin fraction, short interspersed nuclear element (SINE) repeat fraction, and fragile site overlap (**Fig. 2a**). This resulted in 152 significant bins (q < 0.1) spanning 44 topologically associating domains (TADs) containing both established osteosarcoma oncogenes (*MYC*, *CCNE1*, *CDK4*) and tumor suppressor genes (*TP53*, *RB1*, *ATRX*, *PTEN*).

**Figure 2:**
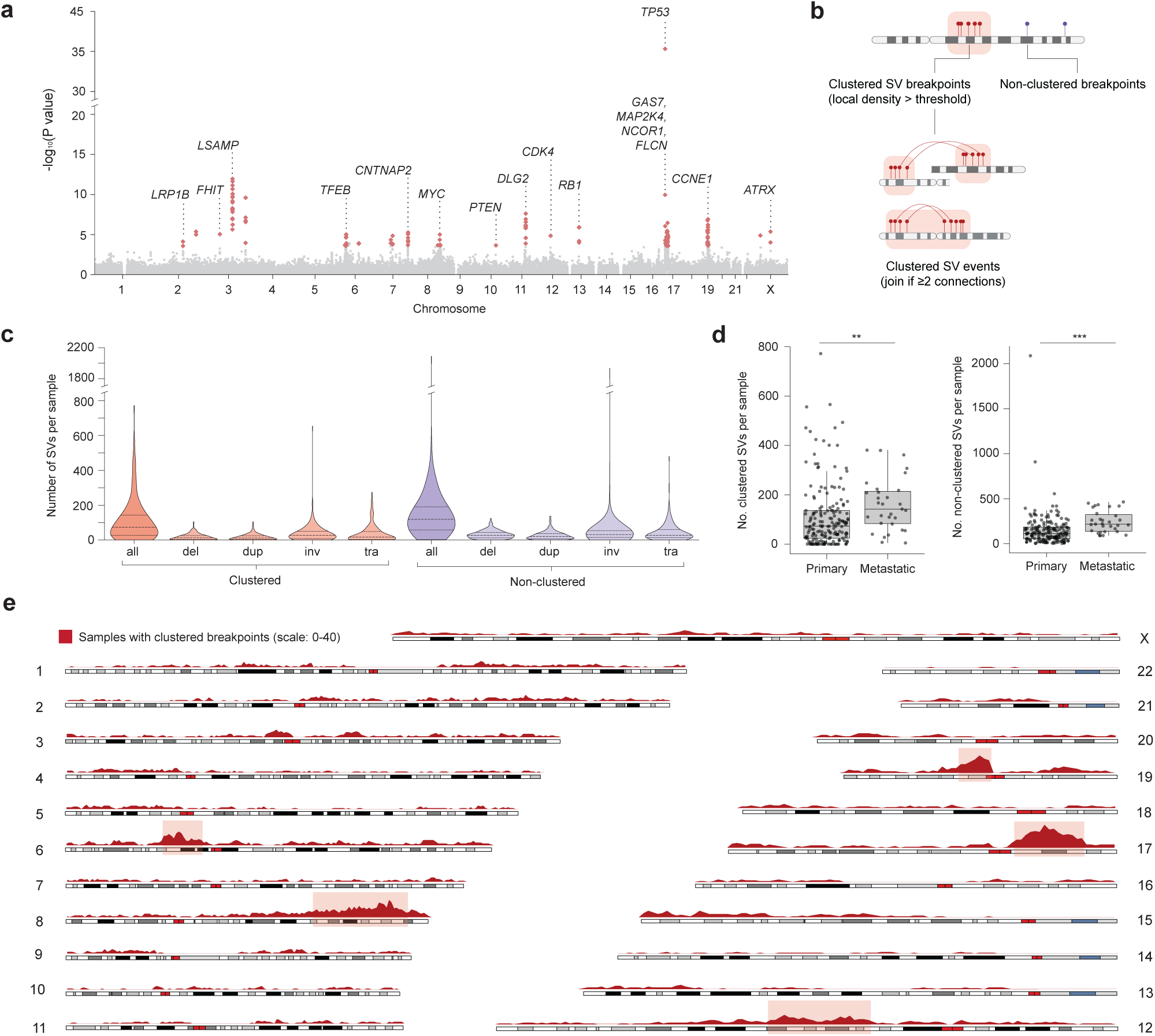
Identification of recurrent SV breakpoints and clustered SV hotspots. **a,** Genome-wide enrichment analysis of SV breakpoints across the cohort (n = 207) using a gamma-Poisson regression model. Each point represents a 50 kb genomic bin; bins significant after Benjamini–Hochberg (BH) correction (*q* < 0.1) are highlighted in red. **b,** Schematic illustrating the categorization of SV breakpoints as clustered or non-clustered. Clustered regions were merged intrachromosomally and linked interchromosomally based on SV connections between them. **c,** Violin plots showing the distribution of clustered and non-clustered SV counts per sample (n = 207), shown overall and stratified by SV type: deletions (del), duplications (dup), inversions (inv), and translocations (tra). **d,** Boxplots comparing clustered (left) and non-clustered (right) SV counts per sample between primary (n = 179) and metastatic (n = 31) tumors. Significance was assessed using a two-sided Mann–Whitney U (MWU) test (**, *P* < 0.01; ***, *P* < 0.001). **e,** Karyoplot showing the number of samples (n = 207) with clustered breakpoints in each 1 Mb window across the genome. The five hotspot clustered SV regions identified are highlighted.

Among these breakpoint-enriched regions, we observed that they were characterized either by simple focal SVs or by dense clusters of rearrangements, which may arise from singular catastrophic events such as chromothripsis or the progressive accumulation of independent rearrangements (**Extended Data Fig. 1c**). To distinguish between these patterns, we classified SV breakpoints for each sample as clustered or non-clustered based on local breakpoint density, an approach adapted from prior rearrangement signature frameworks^18,19^ (**Fig. 2b, Supplementary Table 2**). Clustered SVs (median 74 per sample), defined as SVs with at least one clustered breakpoint, accounted for a substantial proportion of the total SV burden, and the distribution of SV classes (deletions, duplications, inversions, translocations) was similar between clustered and non-clustered SVs (**Fig. 2c**). Metastatic samples had significantly higher burdens of both clustered and non-clustered SVs than primary tumor samples (Wilcoxon rank-sum test, *P* = 2.04 x 10^-3^ and *P* = 3.93 x 10^-6^ respectively; **Fig. 2d**).

We next merged clustered breakpoint regions connected by rearrangements into discrete clustered SV events, identifying 589 events across the cohort, including 142 (24%) spanning multiple chromosomes. At least one clustered SV event was identified in 87% of samples (181/207), with a median of 2 events per sample (IQR: 1–4). To identify regions recurrently affected by these events, we quantified the number of samples with clustered breakpoints across 1 Mb bins genome-wide and merged adjacent high-density bins. This revealed five hotspot loci: 6p21.2–p12.1, 8q22.2–q24.23, 12q13.13–q21.2, 19q11–q13.12, and 17p13.1–11.2. Four of these encompassed established osteosarcoma oncogenes (*CCND3*, *MYC, CDK4/MDM2, CCNE1*) and the fifth fell largely upstream (5’) of *TP53* (**Fig. 2e, Supplementary Table 2**). Overall, 58% of samples (119/207) harbored at least one of these hotspot clustered SV events.

### Hotspot clustered SVs relate to other genomic lesions and show coordinated patterns of co-occurrence and mutual exclusivity

We next sought to characterize hotspot clustered SV events in the context of other genomic lesions in osteosarcoma. Consistent with the known role of complex rearrangements in driving oncogene amplification^20^, we found that samples with clustered SV events at the oncogene-associated hotspot loci showed broad copy number gains spanning several megabases, peaking at the corresponding oncogene (**Fig. 3a**). Samples with the *TP53* upstream clustered SV event, however, showed copy-number loss at and downstream of the *TP53* locus, alongside amplification of the upstream region.

**Figure 3:**
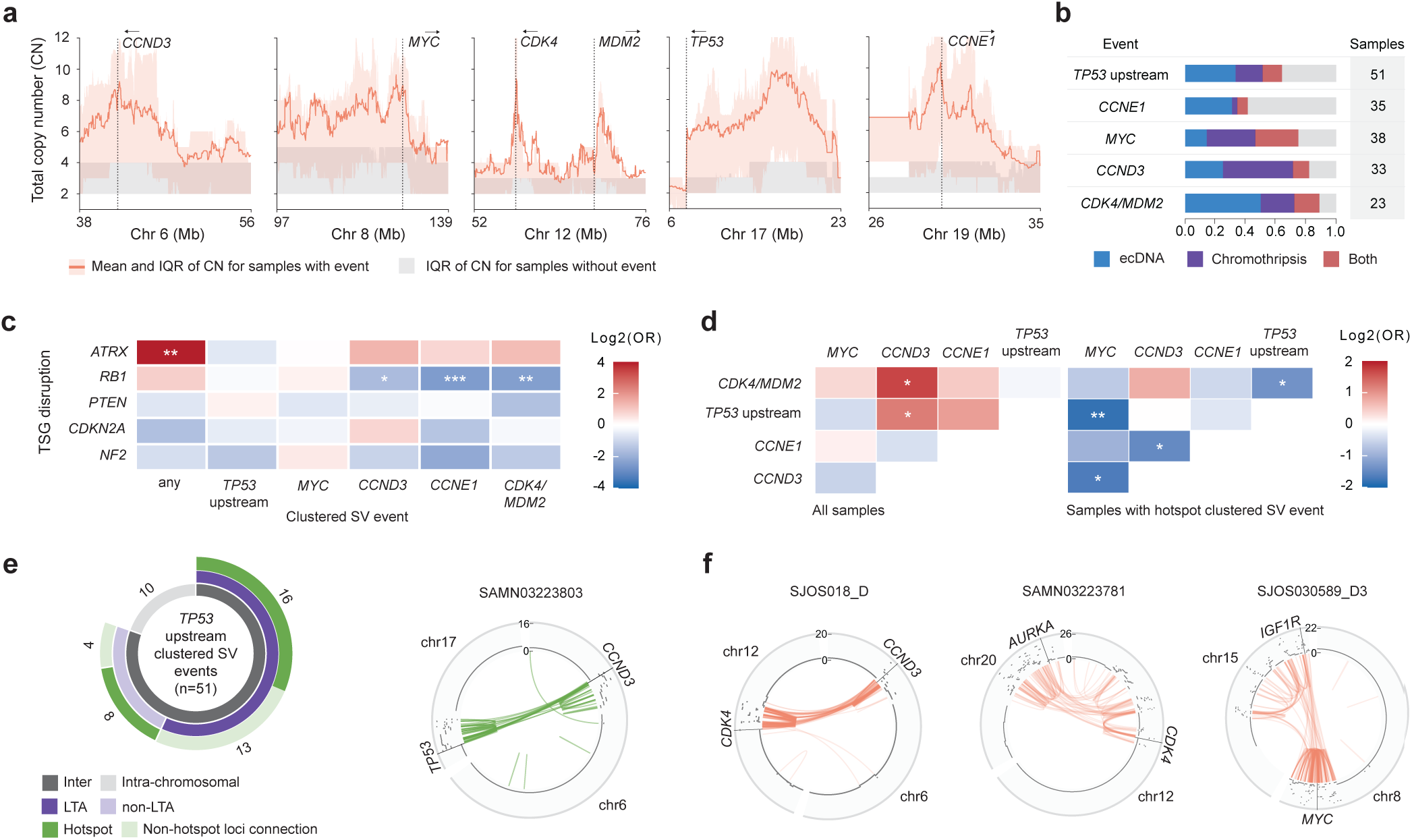
Characterization of hotspot clustered SVs and their associations with other genomic lesions. **a,** Aggregate copy number profiles across hotspot regions. Orange line and shading indicate the mean and interquartile range of copy number for samples with the clustered SV event; gray shading indicates the interquartile range for all other samples. Arrows above gene names indicate the 5′-to-3′ orientation of each gene. Dotted black lines indicate the location of the transcriptional start site of the corresponding gene. **b,** For each hotspot, the proportion of events associated with chromothripsis, ecDNA, both, or neither (bar graph), and the total number of unique-patient samples with the event. **c**, Co-occurrence and mutual exclusivity of tumor suppressor gene (TSG) inactivation and hotspot clustered SV events across samples (n = 203). Associations were assessed using a purity-adjusted Firth logistic regression model, with significance determined after BH correction (*, *q* < 0.05; **, *q* < 0.01, ***, *q* < 0.001). Firth-corrected log_2_ odds ratios (OR) are depicted in the heatmap. **d,** Co-occurrence and mutual exclusivity of hotspot clustered SV events across the entire cohort (n = 203, left) and among samples harboring at least one hotspot event (n = 115, right). Associations were assessed as in c. **e,** Donut chart (left) of *TP53* upstream clustered SV events (n = 51). The inner ring partitions events by whether they involved one or multiple chromosomes, the middle ring by LTA status, and the outer ring by whether the event involved connections to oncogene-associated hotspot loci (i.e., *CCND3*, *CCNE1*, *MYC*, and *CDK4/MDM2*). The middle and outer rings represent subsets of interchromosomal events. Values labeling outer segments indicate the number of events in each category. Circos plot (right) shows a representative LTA event spanning the *TP53* upstream and *CCND3* loci. Arcs indicate SVs; the outer track shows total copy number. **f,** Circos plots showing representative examples of oncogene co-amplification in clustered SV events spanning multiple chromosomes.

We further examined the relationship between hotspot clustered SV events and previously described complex rearrangement processes pervasive in osteosarcoma, namely chromothripsis and extrachromosomal DNA (ecDNA) amplification^6,7,21^. We applied ShatterSeek^22^ to identify chromothripsis based on criteria including equal distribution of SV types and copy number oscillations across 2–3 states, and AmpliconArchitect^23^ to reconstruct ecDNA structures. We found that, across the five hotspot loci, 10–61% of clustered SV events were classified as canonical chromothripsis and 36–67% co-localized with ecDNA amplification (**Fig. 3b**). Notably, ecDNA amplification was selectively enriched at clustered SV hotspot loci, where it occurred at 21.6-fold higher frequency than the genome-wide background (Mann-Whitney U test, *P* = 5.39 × 10^-69^; **Extended Data Fig. 1d**). Among oncogenes recurrently amplified on ecDNA (in ≥ 5 samples), 95% (20/21) mapped to hotspot loci; the sole exception was *IGF1R*. Although ecDNA amplification and chromothripsis were frequently associated with these events, 11–59% exhibited neither feature, suggesting the clustered SV definition captured a broader spectrum of complex rearrangement processes.

We next asked whether intrinsic genomic features could explain why clustered SVs recur at these five loci. Genome-wide, clustered SVs were enriched in earlier-replicating, lower GC-content, SINE-enriched, long terminal repeat (LTR)-depleted, and long interspersed nuclear element (LINE)-depleted regions relative to regions without clustered SVs, though effect sizes were small (rank biserial |*r*| < 0.10; **Extended Data Fig. 1e**). However, when we compared the five hotspot loci specifically to all other clustered SV regions, we found no significant differences across these features apart from a modest enrichment in SINE content. This suggested that certain genomic features may create a more permissive background for the formation of clustered SVs generally, but that their recurrence at specific loci was more consistent with positive selection for the resulting oncogenic alterations.

The high SV density within clustered SV regions raised the question of whether the hotspot events frequently disrupt TAD boundaries, which can alter chromatin architecture and dysregulate gene expression^24^. To assess this, we defined boundary-affecting SVs (BA-SVs) as SVs spanning the full width of a TAD boundary, as previously described^25^. Across samples harboring a given hotspot clustered SV event, TADs overlapping the clustered SV region were significantly more likely to be affected by at least one BA-SV compared to TADs outside the region (generalized estimating equations; **Extended Data Fig. 1f**). This enrichment was consistent across all five hotspot loci: *MYC*, *TP53* upstream, and *CCND3* (*P* < 0.001), *CCNE1* (*P* < 0.01), and *CDK4/MDM2* (*P* < 0.05). Thus, TAD boundary disruption is a frequent consequence of these events, with 60–73% of TADs within clustered SV regions impacted by a BA-SV across the five loci.

We next examined the relationship between clustered SV events and somatic tumor suppressor gene inactivation, which can occur through SCNAs, intragenic SV breakpoints, or mutations including SNVs and small insertions or deletions (indels). We first characterized recurrent SNV and indel driver genes in the cohort, taking the union of mutations across each patient’s samples. Consistent with prior reports, *TP53* (20%), *RB1* (7%) and *ATRX* (5%) emerged as the most frequently mutated genes^5^. Although we also identified *PTEN* and *NF2* as significant drivers, their prevalence was notably lower (1%, 3 patients each) (**Extended Data Fig. 1g**). Integrating mutations with SVs and SCNAs in the non-redundant cohort of 207 samples revealed that *TP53*, *RB1*, and *PTEN* were altered in 76.3% (158), 49.3% (102), and 30% (62) of samples, respectively, with biallelic *TP53* inactivation inferred in 63% (130). *ATRX* alterations were found in 23.2% (48) of samples, although this reflected only SVs and SNVs due to the absence of reliable copy number data for chromosome X. Additionally, *CDKN2A* was disrupted in 24.6% (51) of samples, primarily through deletions.

To account for the lower tumor purity observed in samples lacking detected clustered SV events (**Extended Data Fig. 1h**), all subsequent associations were tested using a Firth logistic regression method with adjustment for tumor purity. *ATRX* disruption was observed exclusively in samples with clustered SVs (Firth purity-adjusted OR = 18.47, *q* = 0.0097), potentially reflecting the role of *ATRX* loss in promoting genomic instability (**Fig. 3c**)^26^. Conversely, *RB1* disruption occurred less frequently in samples with *CCNE1* (OR = 0.18, *q* = 0.0009), *CDK4/MDM2* (OR = 0.19, *q* = 0.0097), or *CCND3* (OR = 0.31, *q* = 0.032) clustered SV events, an inverse relationship that likely reflects functional redundancy, as *CCNE1*, *CDK4*, and *CCND3* are essential components of kinase complexes that phosphorylate and inactivate *RB1*^27^.

We next examined pairwise associations between clustered SV hotspot events to determine whether they co-occurred or were mutually exclusive within tumors. We found that *CDK4*/*MDM2* and *CCND3* clustered SVs co-occurred more frequently than expected (OR = 4.36, *q* = 0.029), which may reflect the cooperative function of CDK4 and cyclin D3 as binding partners within the same cell cycle kinase complex^27^ (**Fig. 3d**). In contrast, restricting the analysis to samples with at least one hotspot event revealed a broader pattern of mutual exclusivity. Tumors were less likely than expected to harbor events at both the *TP53* upstream and *MYC* (OR = 0.21, *q* = 0.0027), *CCND3* and *MYC* (OR = 0.24, *q* = 0.020), *CCND3* and *CCNE1* (OR = 0.28, *q* = 0.038), and *TP53* upstream and *CDK4/MDM2* (OR = 0.30, *q* = 0.041) loci. Overall, 58% of samples with a hotspot event (69/119) had exactly one affected hotspot locus. This pattern suggests that a clustered SV event at one hotspot locus may reduce the selective advantage of acquiring others.

Given that *TP53* upstream clustered SV events were associated with terminal loss of 17p, a feature of the recently described LTA chromothripsis, we sought to determine whether these rearrangements were consistent with this mechanism. Among the 51 samples harboring the *TP53* upstream event, 29 (57%) met the LTA criteria, including biallelic *TP53* inactivation, terminal 17p loss, and inter-chromosomal rearrangements to other clustered SV regions (**Fig. 3e**)^7^. Under the LTA model, these cases arise from a double-strand break within or upstream of *TP53* that results in terminal 17p loss, leaving an unprotected chromosome end that fuses with another chromosome to form a dicentric chromosome. Repeated BFB cycles then generate clustered rearrangements and oncogene amplifications across chromosomes. Across LTA chromothripsis cases, 16 (55%) showed rearrangements connecting the *TP53* upstream locus specifically to one or more of the other hotspot loci, most frequently *CCND3* (7 cases), consistent with the significant co-occurrence of these events, followed by *CCNE1* (6), *CDK4* (2), and *MYC* (2). Among the remaining 22 events not meeting all LTA criteria, 12 (55%) spanned multiple chromosomes, of which 8 involved the other hotspot loci.

The genome-wide prevalence of inter-chromosomal clustered SV events prompted us to investigate whether these rearrangements mediated co-amplification of oncogenes across chromosomes. Across the cohort of 207 samples, 9 had clustered SVs at both the *CCND3* and *CDK4/MDM2* loci, consistent with their observed co-occurrence. In 6 of these cases, a single event jointly affected both loci, including 4 cases with co-amplification of these oncogenes on ecDNA (**Fig. 3f**). Similarly, 7 samples had clustered SVs at both *CCNE1* and *MYC*, with 4 representing a single event spanning both loci. Beyond these inter-hotspot pairings, we identified additional co-amplifications with potential functional relevance: *CDK4*/*MDM2* with *AURKA*, implicated in resistance to CDK4/6 inhibitors^28^, and *MYC* with *IGF1R*, consistent with the role of IGF signaling in sustaining *MYC* activation^29^. Collectively, these findings show that inter-chromosomal clustered SV events can mediate both the co-amplification of oncogenes across distinct loci and simultaneous tumor suppressor gene inactivation with oncogene amplification, as exemplified by LTA. This highlights the capacity of complex rearrangements to drive multiple oncogenic hits within a single event.

### Hotspot clustered SVs drive transcriptional dysregulation

To examine the transcriptional consequences of hotspot clustered SV events, we first leveraged matched bulk RNA-seq data available for 120 tumors from 107 unique patients. We performed differential gene expression analysis comparing samples with and without each hotspot event. This revealed a pattern of broad transcriptional upregulation of genes within each hotspot locus in samples harboring the corresponding event, with 14.5–52.6% of locus genes significantly upregulated compared to 0.1–1.0% of genes elsewhere in the genome (Fisher’s exact test, *q* < 1×10^-18^ for all loci). This included significant upregulation of the implicated oncogenes at the loci (*MYC*, *CCNE1*, *CCND3*, and *CDK4*), consistent with their copy number amplification (**Fig. 4a**). *MDM2*, however, was not differentially expressed among samples with the *CDK4*/*MDM2* event (log_2_FC = 0.33, *q* = 0.50), supporting *CDK4* as the primary oncogenic target at this locus. At the *CCND3* locus, *RUNX2*, *VEGFA*, and *CDC5L* have also been proposed as candidate oncogenes in osteosarcoma^30^. *RUNX2* and *CDC5L* were significantly upregulated (log_2_FC = 0.92, *q* = 0.0013 and log_2_FC = 0.94, *q* = 9.97 x 10^-5^, respectively), although to a lesser extent than *CCND3* (log_2_FC = 2.07, *q* = 5.13 x 10^-11^). *VEGFA* did not reach transcriptome-wide significance (log_2_FC = 0.85, *q* = 0.18). Together, these findings indicate that hotspot clustered SV events are associated with transcriptional upregulation of both the target oncogenes and neighboring genes across each locus.

**Figure 4:**
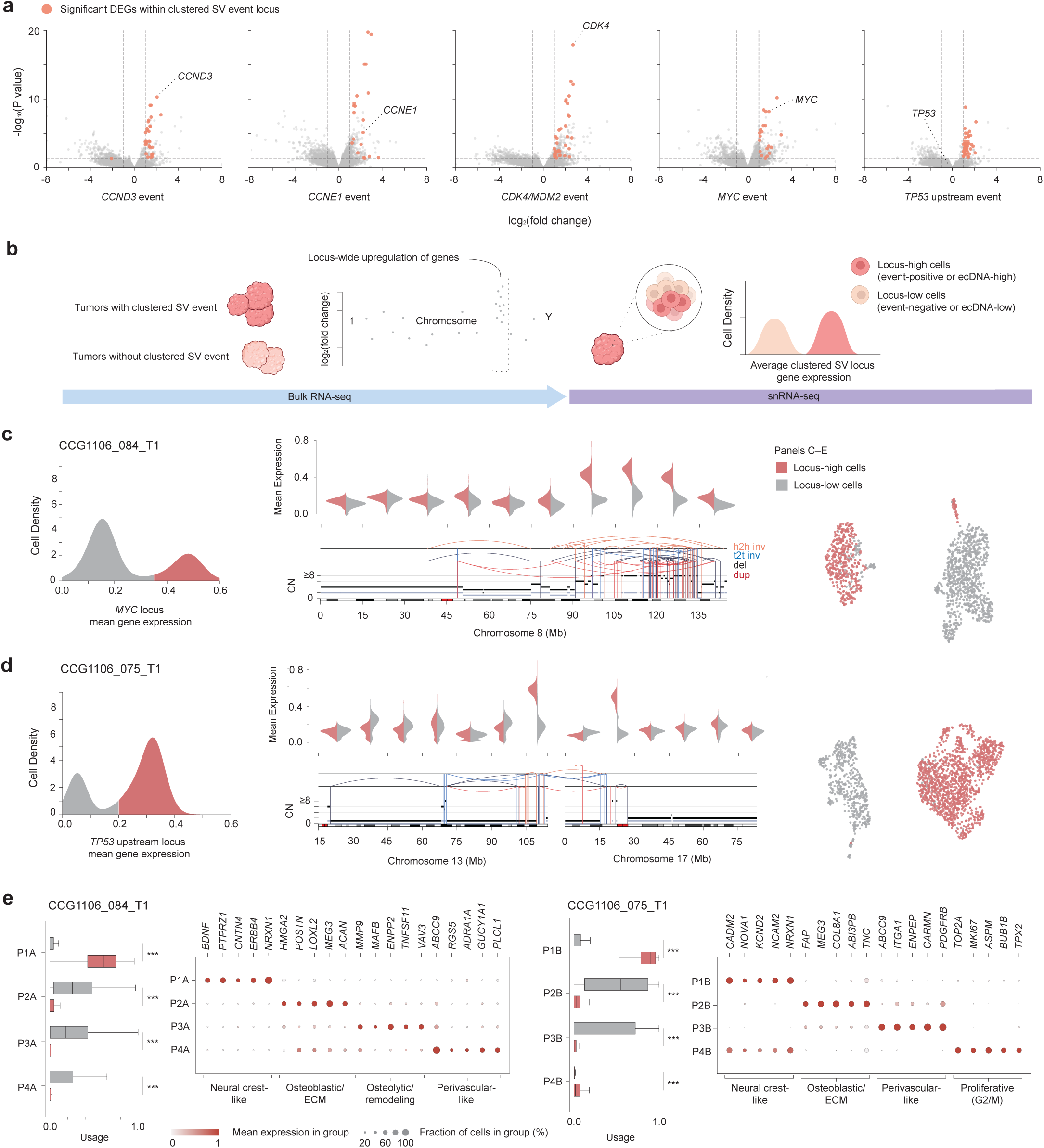
Transcriptional consequences of hotspot clustered SVs. **a,** Volcano plots of differentially expressed genes (DEGs) from bulk RNA-seq in samples with versus without each hotspot event. Significant DEGs within the corresponding hotspot locus are highlighted in orange. **b,** Schematic showing the strategy used to identify event-associated signal in snRNA-seq data, leveraging the locus-wide transcriptional upregulation observed in bulk RNA-seq. **c,** Average expression of genes within the *MYC* hotspot locus across single cells in sample CCG1106_084_T1 (left), showing a bimodal distribution that defines locus-high (n = 410) and locus-low (n = 863) subpopulations. Regional expression scores for the two subpopulations across 15-Mb windows spanning chromosome 8, shown as split violin plots (middle, top). The corresponding genomic landscape, including SVs and CNVs, is shown below, with total and minor allele copy-number in black and gray, respectively (middle, bottom). UMAP colored by inferred event status (right). **d,** As in c, for sample CCG1106_075_T1 harboring a *TP53* upstream clustered SV event, with locus-high (n = 1382) and locus-low (n = 573) cell subpopulations **e,** GEP usage scores for locus-high and locus-low cells in samples CCG1106_084_T1 and CCG1106_075_T1, shown as boxplots for each program (left). Significance was assessed using the two-sided MWU (***, *P* < 0.001). Top contributing genes for each program are shown as a dot plot (right).

Next, we aimed to characterize the transcriptional impact of hotspot clustered SV events at single-cell resolution using matched snRNA-seq data available for 14 samples from 14 unique patients. We identified malignant osteosarcoma cells by annotating cell types using a supervised non-negative matrix factorization (NMF) approach and selecting mesenchymal cells with significant inferred autosomal copy number alterations relative to a diploid reference cell population (**Extended Data Fig. 2a**). To resolve hotspot clustered SV events at the cellular level, we leveraged the locus-wide gene upregulation observed in bulk RNA-seq. For each malignant cell, we computed a regional expression score as the mean expression of genes within the clustered SV locus. In tumors harboring the hotspot event, cells with an elevated score relative to the overall distribution were inferred to carry the alteration and its associated locus-wide transcriptional upregulation (**Fig. 4b**). This revealed two samples with bimodal score distributions, suggesting distinct malignant cell subpopulations: CCG1106_084_T1 (*MYC* event) and CCG1106_075_T1 (*TP53* upstream event) (**Extended Data Fig. 2b**). We designated the high- and low-scoring populations locus-high and locus-low, respectively. This bimodality may reflect either subpopulations with and without the hotspot event or, given the ecDNA-based amplification of these loci, ecDNA-low and ecDNA-high subpopulations due to unequal ecDNA segregation during mitosis.

To confirm that elevated scores specifically reflected effects of the hotspot event rather than global transcriptional or technical variation in these samples, we verified that the expression difference between locus-high and locus-low cells was focal to the event region, with no comparable shifts observed across 15 Mb regions surrounding the locus and genome-wide (**Fig. 4c**, **Fig. 4d**). For CCG1106_075_T1, which had an inter-chromosomal event spanning the *TP53* upstream region and a region on chromosome 13, cells assigned to the locus-high subpopulation showed elevated expression at both loci, further confirming that the score captured event-associated signal.

In both samples, locus-high and locus-low cells clustered separately, reflecting transcriptional heterogeneity between the two subpopulations. Using CytoTRACE2^31^ as a lineage-agnostic measure of differentiation, locus-high cells were the least differentiated subpopulation in both tumors (**Extended Data Fig. 2c**). To characterize these transcriptional differences, we identified gene expression programs (GEPs) in each sample using consensus NMF, retaining four programs per sample (**Supplementary Table 3 and 4**). This unsupervised approach recapitulated the observed clustering, with locus-high and locus-low cells largely confined to distinct programs. Specifically, locus-high cells concentrated within a single program (P1A and P1B in the *MYC* and *TP53* upstream hotspot samples, respectively), except for a minor proliferative subpopulation in the *TP53* upstream sample marked by P4B. Conversely, locus-low cells exhibited greater transcriptional heterogeneity, spanning three programs (P2A–P4A) in the *MYC* sample and two (P2B, P3B) in the *TP53* upstream sample (**Fig. 4e, Extended Data Fig. 2d**).

Despite arising in the context of distinct clustered SV events, the predominant locus-high programs in both samples (P1A and P1B) converged on a neural crest-like state with co-active osteoblastic lineage features (**Extended Data Fig. 2e, f**). Both programs expressed neural markers, including the synaptic adhesion molecule *NRXN1*^32^ and neural cell adhesion molecule *NCAM2*^33^. A pySCENIC analysis of these programs showed transcription factor activity associated with early developmental plasticity^34,35^ (*PITX2* in both, *LIN28B* in P1A) and TGF-beta signaling^36^ (*SMAD3* in both), alongside osteoblastic regulatory activity^37,38^ (*SATB2* in both, *SP7* in P1A). Notably, both programs also showed activity of regulons associated with neural crest specification, including *PAX7* and *SOX9* in P1A and *SOX10* in P1B^39–42^. In contrast, locus-low cells were enriched in more differentiated programs reflecting canonical osteoblastic and perivascular-like states. Together, these findings indicated that cells harboring hotspot clustered SV events and their associated upregulation occupied a more primitive, transcriptionally plastic state diverging from a purely osteoblastic identity, a finding that warrants validation in larger cohorts.

A third sample (SJOS001107_M2) showed uniformly elevated *CCND3* regional expression scores across all malignant cells relative to the cohort-wide distribution, consistent with a homogeneous event-carrying population showing uniform locus-wide upregulation (**Extended Data Fig. 2g**). Supporting this, the *CCND3* clustered SV represented a truncal event, detected across all patient-matched samples including the diagnostic tumor (SJOS001107_D1) and an independent metastatic resection (SJOS001107_M1), indicating early acquisition during tumor evolution. Notably, this patient also had a truncal *CDKN2A/B* deletion, suggesting compounding disruption of cell cycle regulation through early genomic events. In line with the ubiquity of the event, CytoTRACE2 scores across all cells in this sample were comparable to those seen for locus-high cells in samples CCG1106_075_T1 and CCG1106_084_T1, further supporting an association between hotspot clustered SV events and less differentiated cell states.

### Hotspot clustered SVs are often acquired early and continue to evolve over time

Due to the recurrence of clustered SVs at the five hotspot loci, we hypothesized that these events undergo positive selection and represent early drivers of tumor evolution. To infer their phylogenetic timing, we analyzed multi-region and longitudinal samples from 20 patients (mean 2.4 samples per patient), encompassing paired diagnostic, recurrent, and metastatic specimens. We classified SVs as truncal (present in all samples from a patient), shared (present in ≥2 but not all samples), and private (unique to a single sample), as previously described^7^. We found that clustered SVs in hotspot regions were more likely to be truncal compared to those within non-hotspot regions (40.7% vs 30.1%, unadjusted; mixed-effects logistic regression, OR = 1.32, 95% CI [1.15-1.51], *P* = 5.8×10^-5^; **Fig. 5a**), suggesting that hotspot clustered SV events are more frequently acquired early and maintained across tumor evolution.

**Figure 5:**
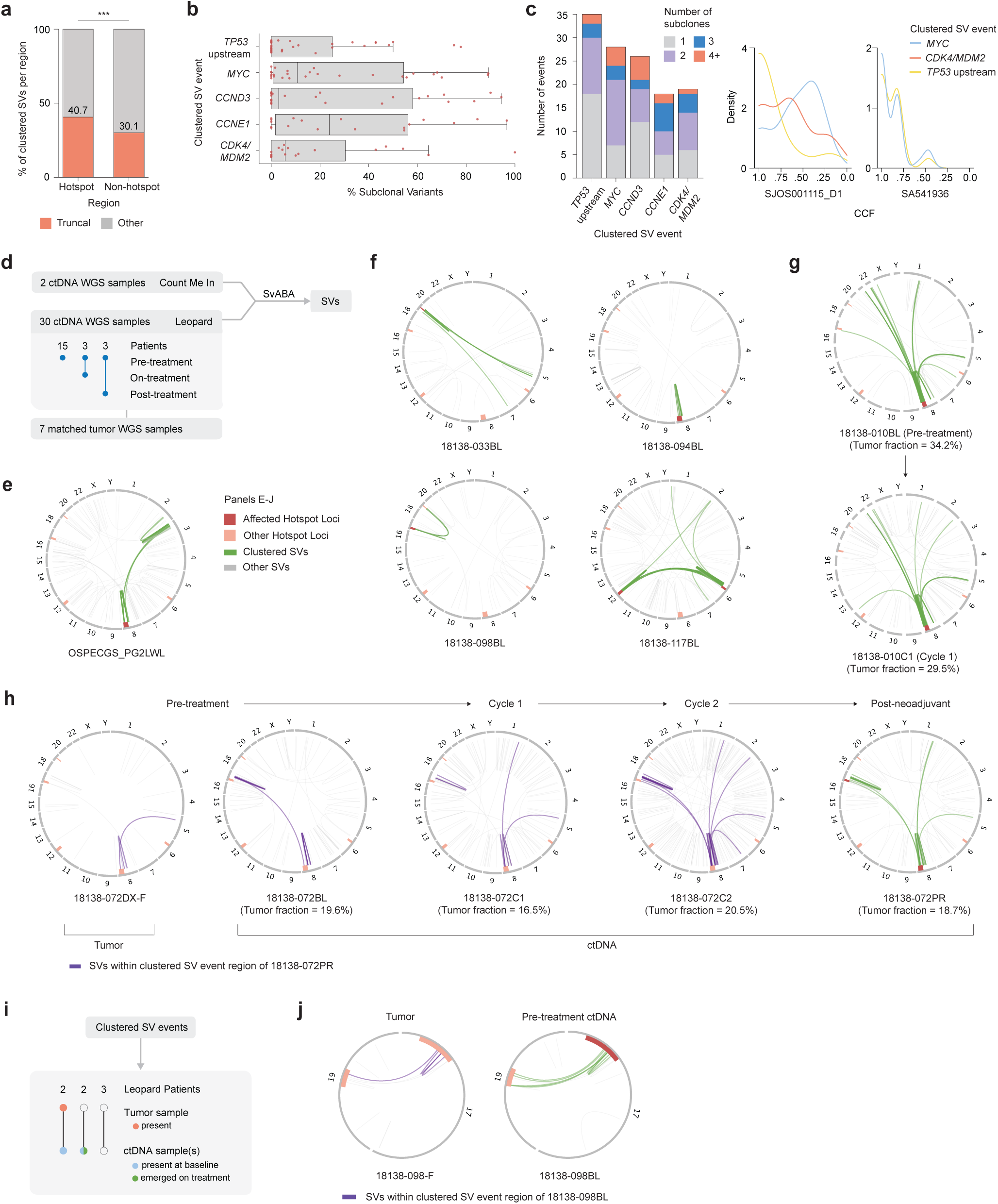
Clonal evolution of hotspot clustered SVs and detection in ctDNA. **a,** Barplot showing percentage of clustered SVs in hotspot (left) and non-hotspot (right) regions classified as truncal or other (shared and private), across 20 patients with multiple longitudinally or spatially distinct samples (47 samples total). Significance from mixed-effects logistic regression with a patient random effect (***P < 0.001). **b,** Box plots of the percentage of subclonal SVs within each clustered SV event identified across the cohort, shown separately for each of the five hotspot loci. Each event is represented by a red dot (n = 35, 28, 26, 18, and 19 for *TP53* upstream, *MYC*, *CCND3*, *CCNE1*, and *CDK4/MDM2*, respectively). **c,** Bar plot (left) showing the number of events displaying 1 (clonal), 2, 3, or ≥4 distinct SV CCF clusters for each hotspot locus. Kernel density estimate (KDE) plots (right) show the distribution of SV CCFs within hotspot events in representative tumors (SJOS001115_D1 and SA541936). **d,** Overview of ctDNA cohorts and analysis workflow. **e,** Circos plot of a clustered SV event identified in a ctDNA sample from the Count Me In (CMI) cohort. Green arcs indicate SVs belonging to a clustered SV event; the red bands mark the affected hotspot loci and orange bands the remaining hotspot loci. **f,** Clustered SV events detected in pre-treatment ctDNA from four Leopard cohort patients. **g,** A clustered SV event in one Leopard cohort patient that persisted from pre-treatment (top) through cycle 1 of treatment (bottom). **h,** One LEOPARD cohort patient’s ctDNA sampled at four timepoints (left to right: pre-treatment, cycle 1, cycle 2, and post-neoadjuvant). A clustered SV event was detected only in the post-neoadjuvant sample (green). At earlier timepoints, SVs mapping to the same loci are shown in purple. **i,** Overview of clustered SV detection across LEOPARD cohort patients comparing matched tumor and ctDNA samples. Filled circles (orange in tumor, blue or green in ctDNA) indicate presence of a clustered SV, open circles indicate no detection. In the ctDNA row, circle color denotes treatment timepoint: blue, present at baseline; green, emerged during treatment. Where multiple ctDNA samples span different timepoints, the circle is split and each segment is colored by its timepoint. **j,** Circos plots of a patient’s matched tumor biopsy (left) and pre-treatment ctDNA (right). A clustered SV event (green) affecting the *TP53* upstream locus was identified in the pre-treatment ctDNA. SVs at the same loci in the tumor sample (purple) did not meet the criteria for a clustered SV.

To further characterize the clonality of hotspot clustered SV events, we used SVClone^43^ to infer the cancer cell fraction (CCF) of individual SVs and quantified the proportion of subclonal SVs (CCF < 0.9) within each hotspot event. While several hotspot events consisted exclusively of clonal SVs (CCF = 1.0), the majority (61.9%) showed varying proportions of subclonal SVs (**Fig. 5b**). The presence of SVs at multiple CCFs within the same event suggested that hotspot loci may undergo recurrent structural evolution, with new rearrangements accumulating over time. To quantify this, we counted the number of CCF clusters within each hotspot event, finding that 49–75% exhibited two or more clusters, suggesting multi-clonal architectures (**Fig. 5c**). At the individual sample level, this pattern was reflected by multimodal distributions of SV CCFs. Together, these analyses indicate that hotspot clustered SV events are established early in tumor development yet continue to evolve, with new rearrangements arising subclonally as tumors progress.

Supporting this notion, we observed evidence of continued selective pressure at the *MYC* locus in a patient with one primary and two metastatic samples, whose truncal *CCND3* event was discussed in the single-cell analysis above. In addition to the *CCND3* event, all three specimens shared a truncal 8q arm-level gain that amplified *MYC* to a copy number of 8. However, a *MYC* clustered SV was detected exclusively in the primary tumor (**Extended Data Fig. 3a**). This event likely arose in the primary tumor after the metastatic lineage had been seeded, further amplifying *MYC* to 14 copies within a detected ecDNA amplicon. These findings suggest that hotspot loci remain under selective pressure for additional structural amplification through clustered SVs, even in the context of pre-existing high-level oncogene copy number gain.

### Liquid biopsies enable non-invasive tracking of ongoing structural evolution

We next hypothesized that circulating tumor DNA (ctDNA) could provide a non-invasive means of following the structural evolution of osteosarcoma genomes over time. To evaluate this, we first analyzed a convenience cohort from the Count Me In (CMI) study^44^ comprising 10 ctDNA WGS samples from 10 patients to determine whether clustered SV events could be resolved. Two of these samples exceeded the 5% tumor fraction threshold we applied for reliable somatic SV detection (**Fig. 5d, Supplementary Table 5**). SV calling with SvABA^16^ identified a clustered SV event involving the *MYC* locus in one case (**Fig. 5e**). We also identified *TP53*-disrupting rearrangements in both cases, demonstrating that clinically relevant SVs can be detected from ctDNA (**Extended Data Fig. 3b**).

We next extended this analysis to the larger Leopard cohort, comprising 30 ctDNA samples with tumor fractions exceeding 5% from 21 patients, all with a pre-treatment baseline specimen^45^. For 5 of these patients, one or more samples were available from subsequent timepoints, spanning early on-treatment, post-neoadjuvant (before local control), and post-treatment draws (**Supplementary Table 5)**. In addition, matched tumor WGS was available for 7 of the 21 patients. In ctDNA, SV calling identified a median of 45 SVs per sample. Five pre-treatment samples harbored clustered SV events, including two at the *MYC* locus, two at *TP53* upstream, and one spanning *CDK4* and *CCND3* (**Fig. 5f, 5g**). A higher proportion of patients with baseline clustered SVs relapsed (3/5, 60%) than those without (6/16, 37.5%), although we were underpowered to assess the significance of this trend. Beyond clustered SV events, SVs affecting key osteosarcoma tumor suppressors across this cohort included rearrangements at *TP53* in three samples, *RB1* in one sample, and *ATRX* in one sample (**Extended Data Fig. 3c**).

The longitudinal samples allowed us to track clustered SV events over the course of treatment. In 3 of the 5 patients, no clustered SV events were present at baseline or at any subsequent timepoint. In a fourth, a clustered SV event at *MYC* detected at baseline persisted in a sample collected early in treatment. The two timepoints differed in several SVs, however, including distinct interchromosomal rearrangements, pointing to ongoing evolution of the locus (**Fig. 5g, Extended Data Fig. 3d**). In the remaining patient, an event spanning the *MYC* and *TP53* upstream loci was detected in a sample taken at the end of neoadjuvant chemotherapy. The matched pre-treatment sample and the intervening neoadjuvant samples during cycle 1 and cycle 2 contained SVs at these loci, though an insufficient number to define a clustered event. Since all four ctDNA samples had similar tumor fractions, the differences in SV counts were unlikely to reflect a purity artifact. The patient’s diagnostic tumor tissue sample likewise lacked a clustered SV event at these loci. The rearrangements therefore appeared to accumulate across successive timepoints, most markedly between the cycle 1 and cycle 2 samples, ultimately reaching the threshold for a clustered SV event in the post-neoadjuvant sample (**Fig. 5h**). Of note, this patient experienced disease progression during treatment after collection of the post-neoadjuvant sample. These findings illustrate the potential of ctDNA to capture ongoing structural evolution at hotspot loci and provide real-time prognostic value as an adjunct to clinical and radiographic assessment during patient trajectories.

Across the 7 Leopard cohort patients with matched tumor WGS, we assessed the concordance of ctDNA-based clustered SV detection with tumor tissue. Every clustered SV event detected in tumor samples was also detected in the corresponding ctDNA (**Fig. 5i**). Results were concordant in 5 patients: 3 had no clustered SVs in either tumor or ctDNA, and 2 harbored the same event in both. These shared events were at the *MYC* locus in one patient (18138-094BL) and spanned the *CDK4* and *CCND3* loci in the other (18138-117BL). In two patients, a clustered SV event was detected in ctDNA but not in the matched tumor, where only simple SVs were seen at the corresponding locus. This included the patient described above with the post-neoadjuvant clustered SV event spanning the *MYC* and *TP53* upstream loci (**Fig. 5h**). In this case, the absence of the event in the tumor sample, which was collected at diagnosis, is consistent with the event having emerged over the course of treatment. In the second patient, a *TP53* upstream clustered SV event was detected in ctDNA at baseline, but only simple SVs were observed at this locus in the matched tumor (**Fig. 5j**). Because these specimens were not necessarily collected contemporaneously, this discordance may reflect spatial or clonal differences in what ctDNA captures relative to a single biopsy, or evolution of the event between collection timepoints.

Across both cohorts, all clustered SV events mapped to the five hotspot loci and were detectable in samples with tumor fractions ranging from 18% to 35%. Together, these results reinforce hotspot clustered SVs as recurrent features of the osteosarcoma genome that can be monitored non-invasively using ctDNA.

## Discussion

The complexity of osteosarcoma genomes, coupled with substantial intertumoral heterogeneity, has presented a long-standing challenge in distinguishing driver events from the background consequences of genomic instability in this aggressive cancer impacting pediatric, adolescent, and adult patients. In this study, we uncovered patterns of complex genomic rearrangements across 236 osteosarcoma tumor-normal paired WGS samples, used matched RNA sequencing to elucidate their transcriptional consequences, and gained new insight into the ongoing evolution and detection of these patterns using ctDNA.

By searching the genome for regions recurrently affected by clustered SV events, we identified five hotspots that encompassed established osteosarcoma oncogenes *CCND3*, *MYC*, *CDK4/MDM2*, and *CCNE1*, and a region largely upstream of *TP53*. Collectively, these hotspot events were present in nearly three-fifths of tumors, indicating that osteosarcoma genomes converge on this limited set of targets despite their otherwise heterogeneous landscapes. We found that oncogene-associated hotspot clustered SV events were associated with broad copy number gains across the locus, peaking at the target oncogenes. Many of these hotspot events were classified as canonical chromothripsis or found to involve ecDNA formation, though they collectively reflected a broader spectrum of complex rearrangement processes. A distinct pattern was seen at the *TP53* upstream locus, which was characterized by copy loss at and downstream of *TP53* together with amplification of sequences upstream of the gene, concordant with the recently reported LTA chromothripsis mechanism^7^.

Three of the five hotspot target genes were direct regulators of the G1/S cell-cycle checkpoint, a process compromised across numerous cancer types^46^. *CCND3*, *CDK4*, and *CCNE1* encode components of cyclin–CDK complexes that phosphorylate and inactivate *RB1*, leading to dysregulated *E2F* activity. Reflecting this functional redundancy, we found that *RB1* inactivation was significantly depleted in tumors harboring *CDK4*, *CCNE1*, or *CCND3* hotspot events. *MYC*, while implicated in widespread cellular processes, also directly activates *E2F* target genes, converging on the same proliferative programs governed by *CCND3*, *CDK4*, and *CCNE1*^47–49^. This functional overlap likely explains the pattern of mutual exclusivity we observed across several hotspot pairs, where one hotspot event reduces the selective pressure to gain a second. Further, the enrichment of early, truncal SVs among these hotspots relative to other clustered SV events suggested that dysregulation of this pathway acts as an early driver of tumorigenesis.

Integrating matched transcriptomic data, we found that hotspot clustered SV events drive the broad upregulation of genes across the affected locus. To characterize the cellular phenotypes associated with these events, we leveraged this locus-wide upregulation to distinguish locus-high cells, inferred to carry the alteration and exhibit its associated expression changes, from locus-low cells lacking this signal. The locus-high cells, identified in two tumors, consistently occupied a more stem-like state and, despite arising from distinct clustered SV events, converged on a neural crest-like transcriptional state. Prior studies have identified neural crest marker expression in chondroprogenitors and an enrichment of neuronal and muscle fate signatures in cancer stem-like cells^50,51^. More broadly, a growing body of evidence across solid tumor types has linked dysregulated differentiation to tumorigenesis, therapeutic resistance, and metastasis^14,52^. Our findings implicate complex genomic rearrangements in this broader pattern.

Previous studies have shown that derivative chromosomes generated by complex genomic rearrangements are primed for ongoing genomic evolution^7^. To explore this phenomenon at clustered SV hotspots, we inferred the CCFs of SVs within each event and found that they spanned a range of values, suggesting that these loci continue to acquire rearrangements over tumor progression. ctDNA derived from blood plasma presents a non-invasive alternative to tumor biopsy that may be particularly useful for longitudinally tracking these changes^53–55^. We showed that clustered SV events are detectable by ctDNA WGS, with all detected events mapping to hotspot loci, further supporting these regions as recurrent targets of complex rearrangement. Importantly, ctDNA captured all clustered SV events detected in matched tumor samples. Serial ctDNA sampling further revealed the evolution of rearrangements at hotspot loci during treatment in two patients. In cases where ctDNA identified events absent from the matched tumor biopsy, this may reflect evolution between sampling timepoints, as well as the limitations of a single biopsy in capturing spatial heterogeneity. These findings highlight the potential of ctDNA for tracking hotspot clustered SV events over the disease course, providing valuable insight into the oncogenic addiction they enable. Such non-invasive monitoring may have diagnostic and prognostic utility in osteosarcoma and other genomically unstable cancers.

While this study provides a comprehensive multi-omic framework for understanding recurrent structural variation in osteosarcoma, several limitations warrant consideration. Fragmented clinical annotations across contributing studies precluded investigation of associations between hotspot events and patient outcomes, limiting our ability to fully establish the clinical relevance of these events. In addition, haplotype phasing of SVs remains challenging with short-read sequencing. Incorporating long-read sequencing would enable haplotype-resolved variant detection, allowing for the reconstruction of allele-specific contexts. Furthermore, the structural evolution of hotspot loci could reflect either discrete, punctuated events or the gradual accumulation of rearrangements, a distinction that will require larger longitudinal or multi-region cohorts to resolve. Finally, whether hotspot clustered SVs drive the adoption of the primitive cell states observed or are enabled by them remains unclear. Addressing this question will require larger cohorts with matched genomic and transcriptomic data, as well as functional studies.

In summary, the identification of clustered SV hotspots in osteosarcoma and the genomic and transcriptomic contexts in which they occur advances our understanding of how complex rearrangements contribute to oncogenesis in this aggressive cancer. More broadly, our findings provide a framework for investigating the patterns, functional consequences, and evolution of complex rearrangements across pediatric and adult cancers. Ultimately, further characterization of these loci and their role in osteosarcoma tumor biology holds the potential to inform clinical monitoring strategies, link these drivers to patient outcomes, and expose new therapeutic vulnerabilities in this disease and others characterized by complex genomes.

## Methods

### Sample selection and whole genome sequencing (WGS) dataset preparation

Osteosarcoma tumor and matched germline WGS data were aggregated from the Therapeutically Applicable Research to Generate Effective Treatments osteosarcoma (TARGET OS; phs000468), International Cancer Genome Consortium (ICGC, EGAC00001000010), St. Jude Cloud (SJC-DS-1001, SJC-DS-1004, SJC-DS-1007, SJC-DS-1008, SJC-DS-1011), Perry et al. (phs000699), and MD Anderson Cancer Center (MDACC; EGAD00001005389) studies. In total, these datasets comprised 260 tumor-normal pairs from 225 patients before quality control.

We additionally performed WGS on 25 frozen tumor tissues and matched normal samples from 17 patients with a diagnosis of osteosarcoma that were seen at Dana-Farber/Boston Children’s Cancer and Blood Disorders Center. Samples were collected with written informed consent and ethics approval by the Dana-Farber Cancer Institute Institutional Review Board under protocol number 17-104. Tumor and normal samples were prepared using PCR-amplified libraries and sequenced on an Illumina NovaSeq6000 to a target depth of 30x. All available clinical characteristics of included patient samples after quality control are summarized in **Supplementary Table 1**.

### Uniform processing of tumor WGS data and somatic variant calling

Raw sequencing reads from all cohorts were mapped to the hg38 build of the human reference genome using BWA-MEM (version 0.7.17)^56^. SNVs and short indels were called using Mutect1 (version 1.1.7)^57^ and Strelka2 (version 2.9.10)^58^. Structural variations (SVs) were called using Manta (version 1.6.0)^15^ and SvABA^16^ (unreleased version; git commit ab39e3f), and we took the union of these calls. SVs were filtered to include those with ≥4 variant reads in tumor, ≤1 in normal, variant allele fraction ≥10%. Allele-specific copy number ratios were determined using HapASeg (version 0.0.5)^59^. Allele-specific absolute copy numbers as well as purity, ploidy, and the number of whole genome doubling (WGD) events were calculated using the ABSOLUTE algorithm (version 1.5)^60^ and involved manual inspection of ABSOLUTE solutions. Loss of heterozygosity (LOH) regions were called when the minor allele copy number was 0. We discarded 17% of samples (49/285) with ABSOLUTE-estimated purity <0.15, including 6 of the 25 newly generated samples. The final dataset comprised 236 samples from 207 participants. Reliable copy-number data for sex chromosomes were not available.

For analyses requiring a non-redundant cohort, we selected one tumor genome per patient, prioritizing primary over metastatic or relapsed samples and, where multiple samples remained, the highest-purity sample.

### Recurrent breakpoint analysis

We divided the genome into 50kb bins, excluding ENCODE blacklist^61^ and assembly gap regions. Following a previously described approach^17^, we used fishHook (version 0.1)^62^ to model a background rate of breakpoints using a Gamma-Poisson model while accounting for genomic covariates. Covariates included replication timing (ENCODE/UW Repli-seq^63^, IMR90 fibroblasts), GC content, mappability^64^, heterochromatin fraction (Roadmap Epigenomics E129 osteoblasts^65^), SINE repeat fraction (RepeatMasker^66^), and overlap with common fragile sites. Genomic annotation tracks were downloaded from the UCSC Genome Browser^67^, with the exception of common fragile site annotations, which were obtained from HumCFS^68^. All covariates were z-score normalized prior to model fitting. We used fishHook *P* values for each bin, representing the probability of observing the observed number of breakpoints or more under the fitted background model, with multiple testing correction performed using the Benjamini-Hochberg procedure. Significant bins (*q* < 0.1) within a topologically associating domain (TAD), defined using a consensus human genome TAD map^69^, were considered one locus and linked to any COSMIC Cancer gene^70^ in that TAD. This analysis was restricted to the non-redundant cohort (n = 207).

### Detection of clustered SVs

Clustered SVs were identified using the bedpeToRearrCatalogue() function from signature.tools.lib (R package)^71^. SVs shorter than 1kb were excluded. A breakpoint was classified as clustered if it belonged to a region with a minimum of 10 breakpoints and an average inter-breakpoint distance less than one-tenth of the genome-wide expected inter-breakpoint distance (calculated as genome size divided by the total number of breakpoints). SVs with at least one clustered breakpoint were classified as clustered SVs. Clustered regions were combined into a single clustered SV event when they were connected by at least two SVs whose breakpoints fell within 10kb of each region. Connected clustered regions on the same chromosome were merged into a single extended locus. To identify hotspot regions across the cohort, the genome was divided into 1Mb windows and the number of samples with clustered breakpoints in each window was counted. Windows in the top 5% by sample count were retained and adjacent windows were merged into contiguous hotspot regions. Clustered SV events were visualized with ReConPlot (version 0.2)^72^ and pyCirclize (version 1.10.1)^73^. KaryoploteR (version 1.24.0)^74^ was used to visualize the number of samples with clustered breakpoints across the genome.

### Detection of chromothripsis

Chromothripsis events were detected using ShatterSeek (version 1.1)^22^. To define an event as chromothripsis it must meet the following criteria: (i) ≥6 interleaved intrachromosomal SVs, (ii) a non-significant fragment joins test, (iii) a significant chromosomal enrichment or exponential breakpoint distribution test, and (iv) copy number oscillation between 2 states across ≥4 contiguous segments. All ShatterSeek *P* values were FDR-corrected. Hotspot clustered SV events were classified as chromothripsis when a ShatterSeek chromothripsis call overlapped the corresponding hotspot locus. Loss-translocation-amplification (LTA) chromothripsis^7^ was identified in samples with a clustered SV at the *TP53* upstream locus and at least one interchromosomal connection to another clustered SV region. LTA calls further required inferred: (i) biallelic *TP53* disruption (≥2 of intragenic SV breakpoint, SNV, and LOH, or a homozygous deletion) and (ii) terminal 17p loss (LOH across ≥75% of 17p through *TP53*, or ≥33% LOH with ≥1Mb at ≥9 copies in that region).

### Detection of extrachromosomal DNA (ecDNA)

To detect ecDNA events, we used the AmpliconSuite pipeline (version 1.5.1), including AmpliconArchitect and AmpliconClassifier^75,76^. Hotspot clustered SV events were inferred to involve ecDNA if an AmpliconClassifier-defined ecDNA amplicon overlapped the locus. The enrichment of ecDNA at hotspot loci was assessed by dividing the genome into non-overlapping 100 kb bins and counting the number of samples with ecDNA amplicons overlapping each bin. Bins were classified as hotspot or non-hotspot based on overlap with hotspot loci, and the distribution of per-bin sample counts was compared between groups using a one-sided Mann-Whitney U test. Fold enrichment was calculated as the ratio of mean sample counts per bin between hotspot and non-hotspot bins.

### Genomic feature enrichment analysis

To assess associations between intrinsic genomic features and clustered SV localization, we divided the genome into 50kb bins and quantified seven genomic features: replication timing, GC content, gene density, heterochromatin fraction, and SINE, LINE, and LTR repeat fractions. Replication timing, GC content, heterochromatin, and repeat annotations were obtained from the same sources described above, while gene density was calculated using GENCODE^77^ annotations. Feature distributions were compared between (i) clustered SV hotspot regions and other clustered SV regions and (ii) all clustered SV regions (including hotspots) and clustered SV void regions using two-sided Mann–Whitney U tests. Effect sizes were quantified using the rank-biserial correlation coefficient (*r*). Ninety-five percent confidence intervals were estimated by bootstrapping (1,000 resamples), and *P* values were adjusted using the Benjamini–Hochberg procedure.

### TAD boundary-affecting SV analysis

TADs were defined as the genomic intervals between consecutive published skeletal muscle TAD boundary calls^78^, excluding TADs overlapping ENCODE blacklisted regions^61^ or assembly gaps. SVs were assigned to a TAD if at least one breakpoint fell within the TAD or its flanking boundaries. Following the PCAWG consortium framework^12^, only intrachromosomal SVs shorter than 2 Mb were considered, and an SV was classified as boundary-affecting (BA-SV) if it spanned the full width of a TAD boundary^25^. Across samples harboring a hotspot clustered SV event, we compared the proportion of TADs impacted by at least one BA-SV between TADs overlapping the clustered SV region and those outside it, excluding TADs with no SVs. Statistical significance was assessed using generalized estimating equations (GEE) with a binomial family, modeling BA-SV presence as the outcome, region (within vs. outside clustered SV locus) as a fixed effect, and sample as the clustering variable to account for within-sample correlation. This analysis was restricted to the non-redundant cohort (n = 207).

### SV clonal evolution analysis

To test whether clustered SVs in hotspot regions were more likely to be truncal while accounting for non-independence of SVs from the same patient, we fit a mixed-effects logistic regression using lme4 (version 1.1.38) with truncal status as the binary outcome, hotspot region status as a fixed effect, and patient as a random intercept. Significance was assessed by likelihood-ratio test against a null model omitting the hotspot term. The model was non-singular, and the effect was robust to leave-one-patient-out analysis. Cancer cell fractions (CCFs) of single-nucleotide variants (SNVs) and SVs were inferred using SVClone^43^ with default parameters and a maximum considered copy number of 10. We limited CCF estimation to the 193 of 207 unique-patient cohort samples that had available BAM files. Because mixed read lengths in several BAM files precluded standard execution of the pipeline, we modified SVClone^43^ to utilize the mean read length across all reads within a given sample. To ensure the reliability of CCF estimates, we further excluded two samples (SJOS030101_D2 and SJOS046149_R1) that exhibited substantial read-length deviations, defined as having either >31% of reads shorter than the dominant read length or a bimodal read-length distribution. Variants were classified as subclonal if their CCF was <0.90.

### SNV driver analysis

For participants with multiple tumor samples, the union of SNVs and indels across all samples was used for this analysis. SNVs and indels in non-coding regions were excluded. De novo signature extraction was performed using SignatureAnalyzer (version 0.0.9)^79^, and extracted signatures were matched to COSMIC v3 reference signatures^80^ by cosine similarity (threshold > 0.85). Signatures identified as artifacts (SBS60) were used to filter the variant set; SNVs and indels with more than 20% attribution to SBS60 were excluded. SNVs and indels were annotated with the Genome Nexus Annotation Pipeline^81^ (GRCh38, MSKCC isoform overrides) to evaluate their potential pathogenic and clinical significance. For SNV driver analysis, mutations were first restricted to those classified by OncoKB (version 7.0)^82,83^ as ‘oncogenic’ or ‘likely oncogenic’, and significantly mutated driver genes were then identified using MutSig2CV (version 3.11)^84^ with default parameters.

### Co-occurrence and mutual exclusivity testing

Associations between tumor suppressor gene (TSG) alterations and clustered SV events, as well as between pairs of clustered SV events, were assessed using purity-adjusted Firth logistic regression. Samples lacking tumor purity estimates (n = 4) were excluded from these analyses. Odds ratios were adjusted for tumor purity, and *P* values were corrected for multiple testing using the Benjamini-Hochberg procedure. TSG alterations were defined as any of the following: an SV breakpoint within the gene body, an exonic SNV or indel classified as ‘oncogenic’ or ‘likely oncogenic’ by OncoKB^82,83^, or copy-number loss (CN < 1.5). For *ATRX*, since no copy-number data were available, SVs classified as deletions that spanned the *ATRX* locus were additionally included as alterations.

### Bulk RNA-seq data generation and processing

Flash-frozen osteosarcoma tissue specimens were submitted to the Broad Institute of MIT and Harvard for RNA extraction and sequencing. Strand-specific RNA sequencing libraries were prepared and sequenced on an Illumina NovaSeq6000 platform, generating 150 bp paired-end reads to a target depth of 50 million read pairs per sample. The sequencing data was obtained in a FASTQ file format, which was then processed using the cumulus implementation of STAR and RSEM to generate count matrices^85–87^. The publicly available bulk RNA-seq data was obtained from TARGET OS (phs000468) and St. Jude’s Cloud in processed, raw counts format, with sequencing and data processing described in the respective manuscripts.

### Differential gene expression analysis using bulk RNA-seq data

Genes with fewer than ten total counts across all samples were excluded from this analysis. Differential expression between tumors with and without a clustered SV at each hotspot region of interest was performed using pyDESeq2^88^ (version 0.5.3), with cohort included as a covariate, as each cohort was processed using different pipelines (design: ∼ cohort + condition). Wald tests were used to assess differential expression, and p-values were adjusted for multiple testing using the Benjamini-Hochberg procedure. Genes with adjusted *P* < 0.05 and absolute log2 fold change > 1 were considered significant. For each hotspot event, the enrichment of upregulated genes within the corresponding locus was assessed by one-sided Fisher’s exact test comparing the proportions of significantly upregulated genes inside and outside the locus. *P* values across the five loci were corrected using the Benjamini-Hochberg procedure.

### Single-nucleus RNA sequencing (snRNA-seq) data processing and malignant cell annotation

Sample sequencing was performed as previously described^14,89^ and generated counts data was obtained from Alex’s Lemonade Stand Foundation single-cell Pediatric Cancer Atlas (accession numbers SCPCP000023 and SCPCP000017). Processing of sequencing data was performed as previously described^89^. Briefly, after initial filtering of low-quality cells, ambient RNA, and potential doublets, iterative leiden clustering using canonical marker genes was performed to annotate cell types, and inferCNV was used to identify and filter for malignant cells per sample.

### Identifying sub-populations of cells with a hotspot clustered SV event

We computed a region expression score for each malignant cell as the mean expression across all genes within each hotspot region, using counts normalized to 10,000 per cell and log1p-transformed. Score distributions were compared between cells from samples with and without the hotspot event (as determined by WGS), to identify a “right-shifted” subpopulation diverging from the bulk distribution. Within samples with the event, individual cells were classified as carrying the alteration and its associated expression changes if their region score exceeded a manually determined threshold based on inspection of the score distribution. To confirm that score-based classification reflected the hotspot event rather than technical or global transcriptional differences, we compared quality-control metrics (detected genes, total counts, mitochondrial fraction, and doublet scores) and region scores at other genomic loci between the classified subpopulations. We used CytoTRACE2^90^ (version 1.1.0.4), a lineage agnostic measure of differentiation based on transcriptional diversity, to infer the degree of cell differentiation. Differences in CytoTRACE2 scores between locus-high and locus-low subpopulations were assessed using two-sided Mann-Whitney U tests.

### Gene expression program analysis

The two samples in which malignant cells exhibited bimodal regional expression scores, suggesting subpopulations with and without the hotspot event, were analyzed using cNMF (version 1.7.0)^91^ to identify cell states. cNMF was run for 100 iterations on 2,000 highly variable genes (HVGs; identified using scanpy (version 1.11.5)) across a range of K values (K = 2–15). The optimal K was selected manually by balancing stability and reconstruction error. Programs that were dominant (normalized usage ≥0.5) in fewer than 5% of cells were excluded as low-usage programs. Programs were functionally annotated by manual review of the top 50 weighted genes against published marker gene sets^92^. To evaluate cross-sample program similarity, we calculated the Spearman rank correlation of gene spectra scores between each pair of retained programs across HVGs shared by both samples.

### Regulon activity inference

We also ran pySCENIC (version 0.12.0) on these samples to infer the activity of known transcription factors^93^. Each run of the program was performed using the default motif (2022 SCENIC+ human motif collection) and search space input (10 kb around the TSS; 500 bp upstream and 100 bp downstream of the TSS) files provided with the pySCENIC package, and a list of known human transcription factors^93,94^. To associate regulon activity with the cNMF-derived gene expression programs, we first assigned cells to their dominant program if normalized usage was ≥0.6 and the ratio of highest to second-highest usage was ≥1.5. AUCell scores for each regulon were then compared between cells assigned to a given program and all remaining cells using two-sided Mann-Whitney U tests. *P* values were adjusted using the Benjamini–Hochberg procedure within each program, and regulons with FDR < 0.05 were considered significantly associated with that program.

### Circulating tumor DNA (ctDNA) and matched tumor WGS data processing and variant calling

The LEOPARD cohort ctDNA samples were obtained through the prospective, multicenter LEOPARD biology study (NCT06068075). Samples were collected from 12 primary study centers. Blood samples were collected at enrollment and at optional pre-specified timepoints during treatment and surveillance. Tumor biopsies were requested but not required for study participation. Where available, WGS data from these biopsies were included to enable comparison with matched ctDNA samples. The LEOPARD and Count Me In (CMI) studies were approved by the IRB at each study center, with informed consent obtained from all patients or their legal guardians and assent obtained according to age when appropriate.

The tumor fraction for both ctDNA samples and the patient-matched tumor samples were determined using ichorCNA (version 2.0.0)^95^ and samples with tumor fraction <5% were excluded. SVs were called using SvABA (version 1.2.0) with an unrelated reference control sample, as matched normal samples were unavailable^16^. SVs were retained if they had ≥4 variant reads in the tumor, ≤1 variant read in the normal, and a variant allele fraction ≥5%. Variants present in the gnomAD SV database (version 4) at allele frequency >0.1 or recurrent across more than 2 samples from independent participants^96^ were filtered out. For ctDNA samples specifically, variants without split-read support were also filtered out, since discordant-only calls are prone to artifacts arising from the short fragment lengths and low tumor fractions characteristic of ctDNA. Clustered SVs were identified using the same approach as described above.

### Visualization

All graphical schematics used in figures were generated using BioRender [Biorender.com]. All other visualization tools are described in the corresponding Methods sections and the GitHub repository. Box-plot boundaries represent the first and third quartiles (Q1, Q3), with the median indicated by the central line; whiskers extend to the most extreme data point within 1.5x the interquartile range. Violin plots show the kernel density distribution of the data, with the Q1, Q3, and median indicated by overlaid lines or a box plot.

---

**Extended Data Figure 1:**
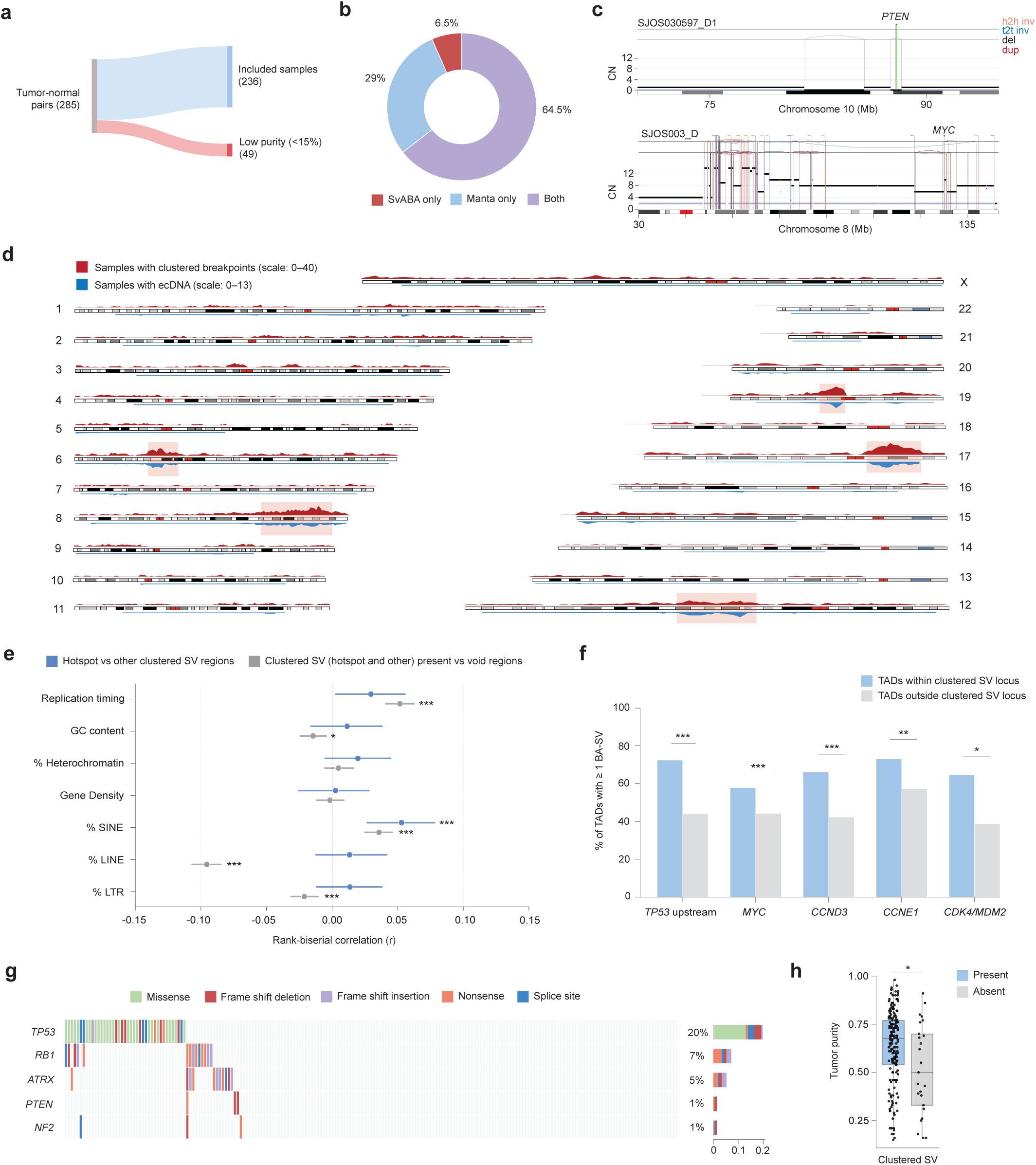
Cohort construction, SV calling quality control, and genomic characterization of clustered SV loci. **a,** Sankey diagram showing sample filtering steps from initial cohort to final analysis set. **b,** Donut chart showing the proportion of the 53,879 total SVs called by Manta only, SvABA only, or both callers. **c,** Rearrangement profiles of representative samples at the *PTEN* locus (SJOS030597_D1, top) and the *MYC* locus (SJOS003_D, bottom). The total and minor allele copy-number are represented in black and gray, respectively. **d**, Karyoplot showing the number of samples (n = 207) with clustered breakpoints (top, red) and ecDNA (bottom, blue) in each 1 Mb window across the genome. The five hotspot clustered SV regions identified are highlighted. **e,** Associations between genomic features and clustered SV loci. Points show rank-biserial correlation effect sizes, with lines representing 95% bootstrap confidence intervals, for two comparisons: hotspot versus other clustered SV loci (blue), and all clustered SV loci versus clustered SV-void regions (gray). Positive values indicate higher feature values in the first group of each comparison. Significance was assessed using two-sided MWU with BH correction (*, *q* < 0.05; **, *q* < 0.01; ***, *q* < 0.001). **f,** Bar plots showing the percentage of TADs impacted by a boundary-affecting SV (BA-SV), comparing regions within versus outside the clustered SV locus for each hotspot, among samples harboring the corresponding event. Significance was assessed using generalized estimating equations (GEE) with a binomial family (*, *P* < 0.05; **, *P* < 0.01; ***, *P* < 0.001). **g,** Oncoprint of SNV/indel driver mutations. Each column represents a patient (n = 207); rows show mutations affecting identified drivers, colored by mutation type. Bar plots on the right show mutation frequency per gene. **h,** Boxplots showing the distribution of tumor purity in samples with (n = 209) and without (n = 27) detected clustered SV events (left). Significance was assessed using a two-sided Mann–Whitney *U* test (*, *P* < 0.05).

**Extended Data Figure 2:**
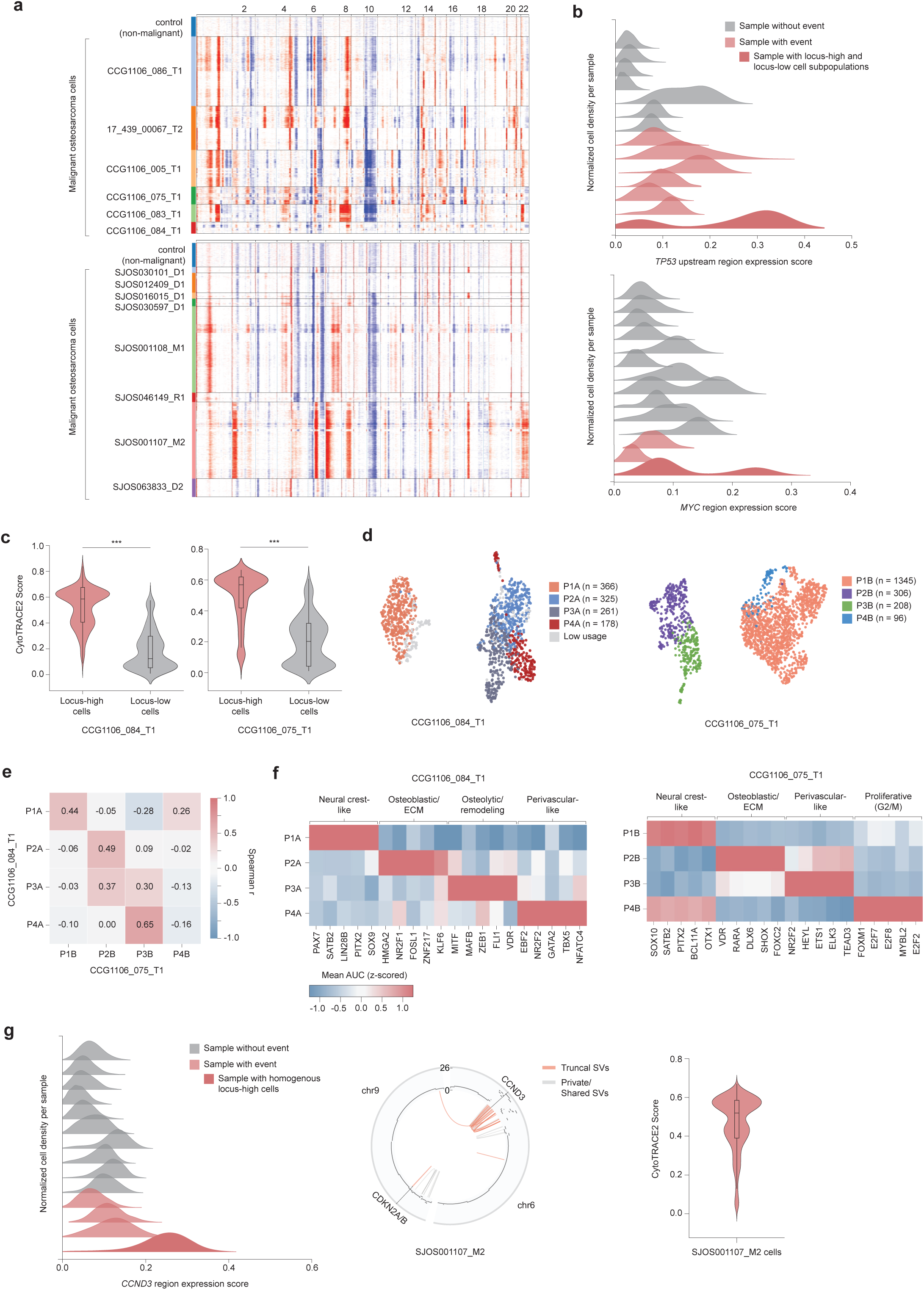
Single-cell transcriptional characterization of clustered SV event-positive cells. **a,** Inferred copy number alterations across autosomes of control non-malignant cells and malignant osteosarcoma cells for 14 snRNA-seq samples from DFCI (top) and St. Jude (bottom) cohorts. **b,** Distribution of regional expression scores for *TP53* upstream (top) and *MYC* (right) loci across cells for each sample, colored by whether the sample harbored the corresponding clustered SV event identified by WGS. **c,** Violin plots showing CytoTRACE2 score distributions (higher scores indicate less differentiated states) by locus status, for samples CCG1106_084_T1 (left) and CCG1106_075_T1 (right). Significance assessed by two-sided MWU (***, *P* < 0.001). **d,** UMAP of cells for the two samples, colored by dominant GEP. **e,** Heatmap showing Spearman correlations (r) between GEPs identified across the two samples. **f,** Heatmap of top regulons for each GEP identified in the two samples, inferred by pySCENIC. **g,** Distribution of *CCND3* regional expression scores across cells from 14 snRNA-seq samples, colored by whether the sample harbored the *CCND3* clustered SV event (left); one sample (SJOS001107_M2) showed uniformly elevated scores. Circos plot of this sample with truncal SVs highlighted in orange (middle). CytoTRACE2 score distribution for all cells (n = 8054) in this sample (right).

**Extended Data Figure 3:**
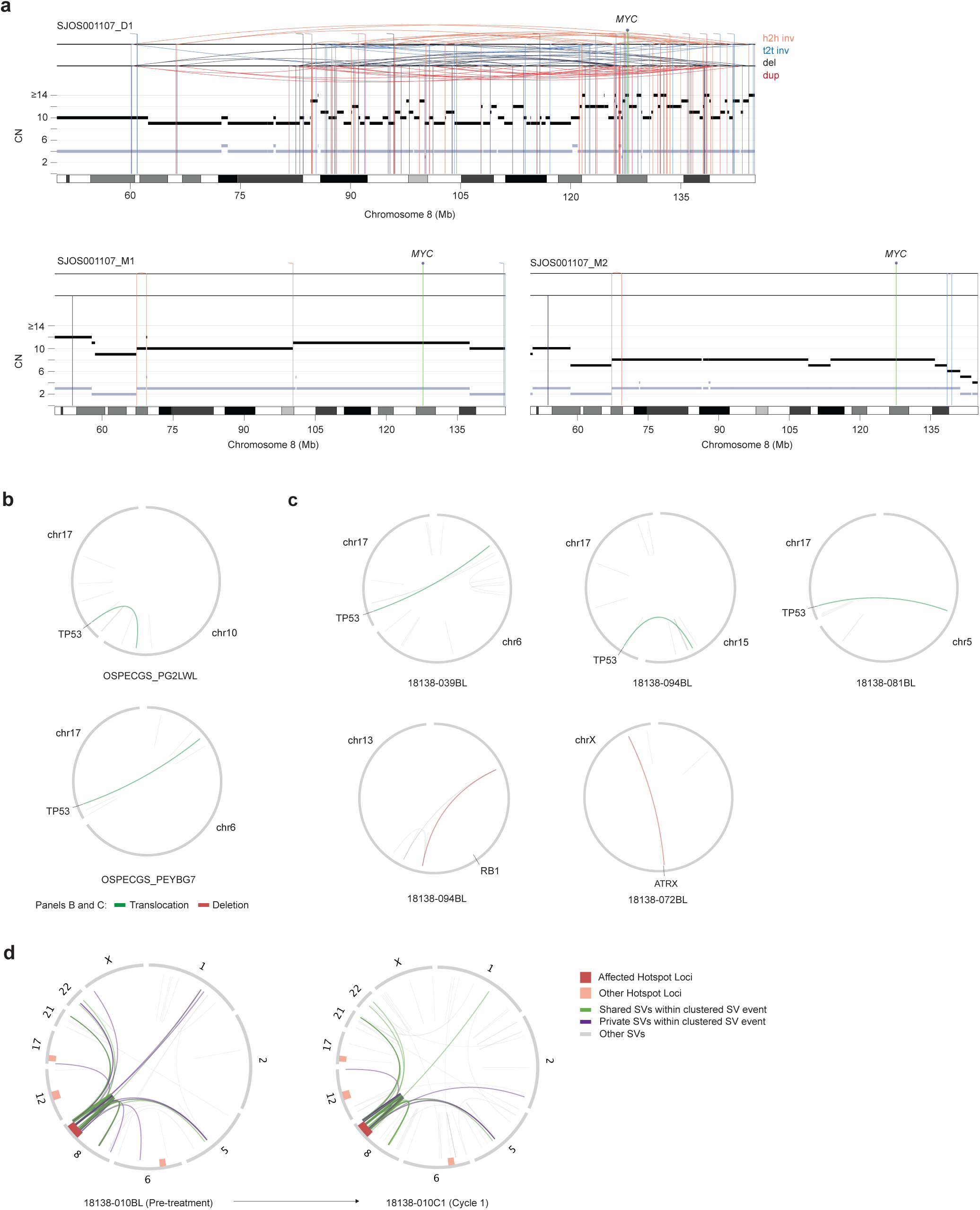
Evolution of clustered SV hotspot loci and ctDNA-based detection of tumor suppressor–disrupting rearrangements. **a,** Rearrangement profiles of samples SJOS001107_D1 (top), SJOS001107_M1 (bottom left), and SJOS001107_M2 (bottom right) at the *MYC* locus. The total and minor allele copy-number are represented in black and gray, respectively. **b**, Circos plots showing TSG-disrupting SVs identified in two CMI cohort samples. **c**, Circos plots showing TSG-disrupting SVs identified in five Leopard cohort samples. **d,** Circos plots of pre-treatment (left) and cycle 1 (right) ctDNA from a patient with a *MYC* clustered SV event detected at both timepoints. SVs shared between timepoints are shown in green; private SVs specific to each sample are shown in purple. Only chromosomes with translocations to the clustered SV locus in either sample are shown.

## Acknowledgements

We gratefully acknowledge the patients, their families and caregivers, and hospital personnel, without whom this research would not have been possible. We also thank Kiara Pontious for her help with project management; Mendy Miller and Saveliy Belkin for their work on reporting and cloud management; all investigators of the LEOPARD and CMI studies; and the other members of the Gillani, Getz, and Crompton Labs at the Dana-Farber Cancer Institute and Broad Institute for their thoughtful comments and feedback throughout this study.

This study makes use of data generated by the St. Jude Children’s Research Hospital including Washington University Pediatric Cancer Genome Project, Genomes for Kids Study and Childhood Solid Tumor Network. The results published here are in part based upon data generated by the Therapeutically Applicable Research to Generate Effective Treatments (https://www.cancer.gov/ccg/research/genome-sequencing/target) initiative, phs000468. The data used for this analysis are available at the Genomic Data Commons (https://portal.gdc.cancer.gov).

This work was supported by Alex’s Lemonade Stand Foundation (ALSF) Young Investigator Awards (YIA) (R.G.); American Society of Clinical Oncology (ASCO) Career Development Award (R.G.); ASCO-Conquer Cancer Sarcoma Foundation of America YIA (R.G.); Boston Children’s Hospital (BCH) Translational Research Program Mentored Translational Investigator Service Award (R.G.); Dana-Farber Cancer Institute (DFCI) Wong Family Award in Translational Oncology (R.G.); DFCI-BCH Pedals for Pediatrics grant (R.G.); Rally Foundation Career Development Award (R.G.); Sarcoma Foundation of America Jay Vernon Jackson Memorial Research Award (R.G.); Department of Defense Peer-Reviewed Cancer Research Program Fellow Career Development Award CA220721 (R.G.); Break Through Cancer’s Defying Osteosarcoma Project (R.G.); Count Me In’s Osteosarcoma Project (K.A.J) and Boehringer Ingelheim Fonds MD Fellowship (E.J.).

This work was additionally supported by NIH grants U2CCA252974 (R.G.); K08CA276701 (R.G.), R37CA244940 (B.D.C) and NIH Project 5U2CCA252974-06 (K.A.J).The content is solely the responsibility of the authors and does not necessarily represent the official views of the National Institutes of Health (NIH).

## Author contributions

Conceptualization: R.G., S.R., T.K., K.A.J., G.G.; Formal analysis: S.R., T.K., D.M., K.P., Y.T., M.T., C.A.R., K.S.K.; Funding acquisition: R.G., G.G., K.A.J., B.D.C., D.S.S., C.Z.Z.; Investigation: all authors; Methodology: S.R., T.K., E.I., B.B., E.J., A.S.; Resources: K.A.J., B.D.C., C.Z.Z, N.B.C., D.M.D., J.J., C.C., L.L.V., E.C., J.M.W.W., K.K., D.S.S.; Supervision: R.G., G.G., K.A.J.; Visualization: S.R.; T.K., Y.T.; Writing –original draft: S.R., T.K., R.G.; Writing –review and editing: all authors

## Data and code availability

The raw sequencing data generated at Dana-Farber Cancer Institute/ Boston Children’s Hospital and the Broad Institute, comprising patient tumor whole-genome sequencing (WGS) with matched normal controls as well as bulk RNA-seq, will be made available at **[dbGaP: phs004090.v2.p1]** (https://dbgap.ncbi.nlm.nih.gov/; NCBI, database of Genotypes and Phenotypes) upon final publication. Whole genome sequencing data for pediatric tumor samples used for analysis in this study were obtained from St. Jude Cloud^10^, dbGaP (phs000468; phs000699) and the European Genome-phenome Archive (EGAD00001005389; EGAD00001002125). The patient tumor snRNA-seq data analyzed in this study are available through the Alex’s Lemonade Stand Foundation, single cell Pediatric Cancer Atlas (ALSF scPCA), accession number SCPCP000017 (https://scpca.alexslemonade.org/projects/SCPCP000017) and SCPCP000023 (https://scpca.alexslemonade.org/projects/SCPCP000023). Of note, a processed version of the bulk patient tumor RNA-seq data is additionally available at SCPCP000017. Sequencing data from ctDNA samples comprising the LEOPARD and CMI cohorts will be made available through dbGaP under their corresponding study projects upon acceptance. Supplementary data and code used in this study’s computational analysis will be publicly available on GitHub at https://github.com/gillanilab/osteosarcoma-wgs-clusteredSVs upon final publication. Any additional information will be made available from the corresponding author, Riaz Gillani, M.D.

## Conflicts of interest

K.A.J reports consulting for Recordati and research funding from AstraZeneca. R.G. has equity in Google, Microsoft, Amazon, Apple, Moderna, Pfizer, and Vertex Pharmaceuticals; his spouse is employed by Carrum Health. The other authors declare that they have no competing interests.

## References

1. Beird, H. C. et al. Osteosarcoma. Nat. Rev. Dis. Primer 8, 77 (2022).

2. Gorlick, R. et al. Children’s Oncology Group’s 2013 blueprint for research: Bone tumors. Pediatr. Blood Cancer 60, 1009–1015 (2013).

3. Ferrari, S. et al. Postrelapse Survival in Osteosarcoma of the Extremities: Prognostic Factors for Long-Term Survival. J. Clin. Oncol. 21, 710–715 (2003).

4. Meltzer, P. S. & Helman, L. J. New Horizons in the Treatment of Osteosarcoma. N. Engl. J. Med. 385, 2066–2076 (2021).

5. Chen, X. et al. Recurrent Somatic Structural Variations Contribute to Tumorigenesis in Pediatric Osteosarcoma. Cell Rep. 7, 104–112 (2014).

6. Greenhalgh, R. et al. The landscape of structural variation in pediatric cancer. Cancer Cell 44, 1029–1044.e6 (2026).

7. Valle-Inclan, J. E. et al. Ongoing chromothripsis underpins osteosarcoma genome complexity and clonal evolution. Cell 188, 352–370.e22 (2025).

8. Behjati, S. et al. Recurrent mutation of IGF signalling genes and distinct patterns of genomic rearrangement in osteosarcoma. Nat. Commun. 8, 15936 (2017).

9. Meijer, D. M. et al. The Variable Genomic Landscape During Osteosarcoma Progression: Insights From a Longitudinal WGS Analysis. https://doi.org/10.1002/gcc.23253 doi:10.1002/gcc.23253.

10. McLeod, C. et al. St. Jude Cloud: A Pediatric Cancer Genomic Data-Sharing Ecosystem. Cancer Discov. 11, 1082–1099 (2021).

11. Perry, J. A. et al. Complementary genomic approaches highlight the PI3K/mTOR pathway as a common vulnerability in osteosarcoma. Proc. Natl. Acad. Sci. U. S. A. 111, E5564–5573 (2014).

12. Aaltonen, L. A. et al. Pan-cancer analysis of whole genomes. Nature 578, 82–93 (2020).

13. Wu, C.-C. et al. Immuno-genomic landscape of osteosarcoma. Nat. Commun. 11, 1008 (2020).

14. Tanaka, Y. et al. Metastatic Osteosarcoma is Characterized by Loss of Osteoblastic Lineage Fidelity. 2026.06.10.731352 Preprint at 10.64898/2026.06.10.731352 (2026).

15. Chen, X., et al. Manta: rapid detection of structural variants and indels for germline and cancer sequencing applications. Bioinformatics 32, 1220–1222 (2016).

16. Wala, J. A. et al. SvABA: genome-wide detection of structural variants and indels by local assembly. Genome Res. https://doi.org/10.1101/gr.221028.117 (2018) doi:10.1101/gr.221028.117.

17. Rheinbay, E. et al. Analyses of non-coding somatic drivers in 2,658 cancer whole genomes. Nature 578, 102–111 (2020).

18. Degasperi, A. et al. A practical framework and online tool for mutational signature analyses show intertissue variation and driver dependencies. Nat. Cancer 1, 249–263 (2020).

19. Nik-Zainal, S. et al. Landscape of somatic mutations in 560 breast cancer whole-genome sequences. Nature 534, 47–54 (2016).

20. Shoshani, O. et al. Chromothripsis drives the evolution of gene amplification in cancer. Nature 591, 137–141 (2021).

21. Bailey, C. et al. Origins and impact of extrachromosomal DNA. Nature 635, 193–200 (2024).

22. Cortés-Ciriano, I. et al. Comprehensive analysis of chromothripsis in 2,658 human cancers using whole-genome sequencing. Nat. Genet. 52, 331–341 (2020).

23. Deshpande, V. et al. Exploring the landscape of focal amplifications in cancer using AmpliconArchitect. Nat. Commun. 10, 392 (2019).

24. Lupiáñez, D. G. et al. Disruptions of Topological Chromatin Domains Cause Pathogenic Rewiring of Gene-Enhancer Interactions. Cell 161, 1012–1025 (2015).

25. Akdemir, K. C. et al. Disruption of chromatin folding domains by somatic genomic rearrangements in human cancer. Nat. Genet. 52, 294–305 (2020).

26. Lovejoy, C. A. et al. Loss of ATRX, Genome Instability, and an Altered DNA Damage Response Are Hallmarks of the Alternative Lengthening of Telomeres Pathway. PLOS Genet. 8, e1002772 (2012).

27. Pellarin, I. et al. Cyclin-dependent protein kinases and cell cycle regulation in biology and disease. Signal Transduct. Target. Ther. 10, 11 (2025).

28. Wander, S. A. et al. The Genomic Landscape of Intrinsic and Acquired Resistance to Cyclin-Dependent Kinase 4/6 Inhibitors in Patients with Hormone Receptor–Positive Metastatic Breast Cancer. Cancer Discov. 10, 1174–1193 (2020).

29. Lee, J.-S. et al. The insulin and IGF signaling pathway sustains breast cancer stem cells by IRS2/PI3K-mediated regulation of MYC. Cell Rep. 41, (2022).

30. Gupta, S. et al. RUNX2 (6p21.1) Amplification in Osteosarcoma. Hum. Pathol. 94, 23–28 (2019).

31. Kang, M. et al. Improved reconstruction of single-cell developmental potential with CytoTRACE 2. Nat. Methods 22, 2258–2263 (2025).

32. Südhof, T. C. Synaptic Neurexin Complexes: A Molecular Code for the Logic of Neural Circuits. Cell 171, 745–769 (2017).

33. Kulahin, N. et al. Structural Model and *trans*-Interaction of the Entire Ectodomain of the Olfactory Cell Adhesion Molecule. Structure 19, 203–211 (2011).

34. Hennchen, M. et al. Lin28B and Let-7 in the Control of Sympathetic Neurogenesis and Neuroblastoma Development. J. Neurosci. Off. J. Soc. Neurosci. 35, 16531–16544 (2015).

35. Hayashi, M., et al. *Pitx2* Prevents Osteoblastic Transdifferentiation of Myoblasts by Bone Morphogenetic Proteins*. J. Biol. Chem. 283, 565–571 (2008).

36. Massagué, J. & Sheppard, D. TGF-β signaling in health and disease. Cell 186, 4007–4037 (2023).

37. Dobreva, G. et al. SATB2 Is a Multifunctional Determinant of Craniofacial Patterning and Osteoblast Differentiation. Cell 125, 971–986 (2006).

38. Hojo, H., Ohba, S., He, X., Lai, L. P. & McMahon, A. P. Sp7/Osterix Is Restricted to Bone-Forming Vertebrates where It Acts as a Dlx Co-factor in Osteoblast Specification. Dev. Cell 37, 238–253 (2016).

39. Cheung, M. & Briscoe, J. Neural crest development is regulated by the transcription factor Sox9. Development 130, 5681–5693 (2003).

40. Kim, J., Lo, L., Dormand, E. & Anderson, D. J. SOX10 Maintains Multipotency and Inhibits Neuronal Differentiation of Neural Crest Stem Cells. Neuron 38, 17–31 (2003).

41. Basch, M. L., Bronner-Fraser, M. & García-Castro, M. I. Specification of the neural crest occurs during gastrulation and requires Pax7. Nature 441, 218–222 (2006).

42. Betters, E., Liu, Y., Kjaeldgaard, A., Sundström, E. & García-Castro, M. I. Analysis of early human neural crest development. Dev. Biol. 344, 578–592 (2010).

43. Cmero, M. et al. Inferring structural variant cancer cell fraction. Nat. Commun. 11, 730 (2020).

44. Wong, J. M. et al. Abstract 630: Directly engaging participants in rare cancer research is feasible: the osteosarcoma and leiomyosarcoma projects. Cancer Res. 85, 630 (2025).

45. Shulman, D. S. et al. Prospective evaluation of pre-treatment ctDNA burden in localized osteosarcoma to identify patients with inferior outcomes: A report from the LEOPARD study. J. Clin. Oncol. 42, 11510–11510 (2024).

46. Knudsen, E. S. et al. Pan-cancer molecular analysis of the RB tumor suppressor pathway. Commun. Biol. 3, 158 (2020).

47. Sears, R., Ohtani, K. & Nevins, J. R. Identification of positively and negatively acting elements regulating expression of the E2F2 gene in response to cell growth signals. Mol. Cell. Biol. 17, 5227–5235 (1997).

48. Adams, M. R., Sears, R., Nuckolls, F., Leone, G. & Nevins, J. R. Complex transcriptional regulatory mechanisms control expression of the E2F3 locus. Mol. Cell. Biol. 20, 3633–3639 (2000).

49. Mateyak, M. K., Obaya, A. J. & Sedivy, J. M. c-Myc Regulates Cyclin D-Cdk4 and -Cdk6 Activity but Affects Cell Cycle Progression at Multiple Independent Points. Mol. Cell. Biol. 19, 4672–4683 (1999).

50. Budhathoki, Y. et al. Integrative Single-cell and Spatial Transcriptomic Analysis of Osteosarcoma Reveals Conserved and Distinct Ecosystems Across Sites and Species. 2026.01.13.698472 Preprint at 10.64898/2026.01.13.698472 (2026).

51. Truong, D. D. et al. Mapping the Single-Cell Differentiation Landscape of Osteosarcoma. Clin. Cancer Res. 30, 3259–3272 (2024).

52. Pérez-González, A., Bévant, K. & Blanpain, C. Cancer cell plasticity during tumor progression, metastasis and response to therapy. Nat. Cancer 4, 1063–1082 (2023).

53. Shulman, D. S. & Crompton, B. D. Emerging Role of Blood-based Biomarkers in Sarcomas. Hematol. Oncol. Clin. North Am. 39, 679–692 (2025).

54. Shah, A. T. et al. A Comprehensive Circulating Tumor DNA Assay for Detection of Translocation and Copy-Number Changes in Pediatric Sarcomas. Mol. Cancer Ther. 20, 2016–2025 (2021).

55. Barris, D. M. et al. Detection of circulating tumor DNA in patients with osteosarcoma. Oncotarget 9, 12695–12704 (2018).

56. Li, H. & Durbin, R. Fast and accurate short read alignment with Burrows–Wheeler transform. Bioinformatics 25, 1754–1760 (2009).

57. Cibulskis, K. et al. Sensitive detection of somatic point mutations in impure and heterogeneous cancer samples. Nat. Biotechnol. 31, 213–219 (2013).

58. Kim, S. et al. Strelka2: fast and accurate calling of germline and somatic variants. Nat. Methods 15, 591–594 (2018).

59. Priebe, O. et al. Haplotype-aware segmentation with HapASeg increases accuracy of detecting homolog-specific somatic copy number alterations. Genome Biol. 27, 83 (2026).

60. Carter, S. L. et al. Absolute quantification of somatic DNA alterations in human cancer. Nat. Biotechnol. 30, 413–421 (2012).

61. Amemiya, H. M., Kundaje, A. & Boyle, A. P. The ENCODE Blacklist: Identification of Problematic Regions of the Genome. Sci. Rep. 9, 9354 (2019).

62. Imielinski, M., Guo, G. & Meyerson, M. Insertions and Deletions Target Lineage-Defining Genes in Human Cancers. Cell 168, 460–472.e14 (2017).

63. Hansen, R. S. et al. Sequencing newly replicated DNA reveals widespread plasticity in human replication timing. Proc. Natl. Acad. Sci. U. S. A. 107, 139–144 (2010).

64. Karimzadeh, M., Ernst, C., Kundaje, A. & Hoffman, M. M. Umap and Bismap: quantifying genome and methylome mappability. Nucleic Acids Res. 46, e120 (2018).

65. Bernstein, B. E. et al. The NIH Roadmap Epigenomics Mapping Consortium. Nat. Biotechnol. 28, 1045–1048 (2010).

66. Smit, A., Hubley, R. & Green, P. RepeatMasker Open-4.0. (2015).

67. Casper, J. et al. The UCSC Genome Browser database: 2026 update. Nucleic Acids Res. 54, D1331–D1335 (2026).

68. Kumar, R. et al. HumCFS: a database of fragile sites in human chromosomes. BMC Genomics 19, 985 (2019).

69. Singh, R. & Berger, B. Deciphering the species-level structure of topologically associating domains. 2021.10.28.466333 Preprint at 10.1101/2021.10.28.466333 (2021).

70. Tate, J. G. et al. COSMIC: the Catalogue Of Somatic Mutations In Cancer. Nucleic Acids Res. 47, D941–D947 (2019).

71. Degasperi, A. et al. A practical framework and online tool for mutational signature analyses show intertissue variation and driver dependencies. Nat. Cancer 1, 249–263 (2020).

72. Espejo Valle-Inclán, J. & Cortés-Ciriano, I. ReConPlot: an R package for the visualization and interpretation of genomic rearrangements. Bioinformatics 39, btad719 (2023).

73. Shimoyama, Y. pyCirclize: Circular visualization in Python. (2022).

74. Gel, B. & Serra, E. karyoploteR: an R/Bioconductor package to plot customizable genomes displaying arbitrary data. Bioinformatics 33, 3088–3090 (2017).

75. Deshpande, V. et al. Exploring the landscape of focal amplifications in cancer using AmpliconArchitect. Nat. Commun. 10, 392 (2019).

76. Luebeck, J. et al. AmpliconSuite: an end-to-end workflow for analyzing focal amplifications in cancer genomes. 2024.05.06.592768 Preprint at 10.1101/2024.05.06.592768 (2024).

77. Mudge, J. M. et al. GENCODE 2025: reference gene annotation for human and mouse. Nucleic Acids Res. 53, D966–D975 (2025).

78. Dunham, I. et al. An integrated encyclopedia of DNA elements in the human genome. Nature 489, 57–74 (2012).

79. Taylor-Weiner, A. et al. Scaling computational genomics to millions of individuals with GPUs. Genome Biol. 20, 228 (2019).

80. Alexandrov, L. B. et al. The repertoire of mutational signatures in human cancer. Nature 578, 94–101 (2020).

81. de Bruijn, I. et al. Genome Nexus: A Comprehensive Resource for the Annotation and Interpretation of Genomic Variants in Cancer. JCO Clin. Cancer Inform. e2100144 (2022) doi:10.1200/CCI.21.00144.

82. Suehnholz, S. P. et al. Quantifying the Expanding Landscape of Clinical Actionability for Patients with Cancer. Cancer Discov. 14, 49–65 (2024).

83. Chakravarty, D. et al. OncoKB: A Precision Oncology Knowledge Base. JCO Precis. Oncol. 1–16 (2017) doi:10.1200/PO.17.00011.

84. Lawrence, M. S. et al. Discovery and saturation analysis of cancer genes across 21 tumour types. Nature 505, 495–501 (2014).

85. Li, B. et al. Cumulus provides cloud-based data analysis for large-scale single-cell and single-nucleus RNA-seq. Nat. Methods 17, 793–798 (2020).

86. Dobin, A. et al. STAR: ultrafast universal RNA-seq aligner. Bioinformatics 29, 15–21 (2013).

87. Li, B. & Dewey, C. N. RSEM: accurate transcript quantification from RNA-Seq data with or without a reference genome. BMC Bioinformatics 12, 323 (2011).

88. Muzellec, B., Teleńczuk, M., Cabeli, V. & Andreux, M. PyDESeq2: a python package for bulk RNA-seq differential expression analysis. Bioinformatics 39, btad547 (2023).

89. Hawkins, A. G. et al. The Single-cell Pediatric Cancer Atlas: Data portal and open-source tools for single-cell transcriptomics of pediatric tumors. 2024.04.19.590243 Preprint at 10.1101/2024.04.19.590243 (2025).

90. Kang, M. et al. Improved reconstruction of single-cell developmental potential with CytoTRACE 2. Nat. Methods 22, 2258–2263 (2025).

91. Kotliar, D. et al. Identifying gene expression programs of cell-type identity and cellular activity with single-cell RNA-Seq. eLife 8, e43803 (2019).

92. Liberzon, A. et al. The Molecular Signatures Database Hallmark Gene Set Collection. Cell Syst. 1, 417–425 (2015).

93. Aibar, S. et al. SCENIC: single-cell regulatory network inference and clustering. Nat. Methods 14, 1083–1086 (2017).

94. Lambert, S. A. et al. The Human Transcription Factors. Cell 172, 650–665 (2018).

95. Adalsteinsson, V. A. et al. Scalable whole-exome sequencing of cell-free DNA reveals high concordance with metastatic tumors. Nat. Commun. 8, 1324 (2017).

96. Guez, J. et al. Integrating 730,947 exome sequences with clinical literature improves gene discovery. 2026.03.23.26349081 Preprint at 10.64898/2026.03.23.26349081 (2026).

